# local adaptation and host specificity to copepod intermediate hosts by the *Schistocephalus solidus* tapeworm

**DOI:** 10.1101/2022.05.15.492025

**Authors:** Kum C. Shim, Christopher Peterson, Daniel I. Bolnick

## Abstract

We investigated if there was local adaptation and host specify in the tapeworm *Schistocephalus solidus* to its copepod first intermediate hosts. The tapeworm is locally adapted and host specific to its threespine stickleback second intermediate host. We exposed copepods from five lakes in Vancouver Island (BC, Canada) to local (i.e. same lake) and foreign tapeworms in a reciprocal exposure experiment. Results indicate that the tapeworm is not locally adapted to the copepods, but there was host specificity as a copepod genus was more parasitized than another genus.

## Introduction

One of the most intriguing features of parasites with complex life cycles is their ability to infect several very disparate hosts during each of their life stages (Schmid-Hempel 2011). Transmission of these parasites (especially helminths) usually involves search for, and penetration of, their intermediate hosts. These parasites then passively infect their final hosts when the intermediate hosts are predated. Thus, complex-life-cycle parasites usually lower the overall fitness of their intermediate hosts (i.e. increased predation), and cannot be as selective on infecting final hosts (Schmid-Hempel 2011, Poulin 2007, Noble et al. 1989). Accordingly, these parasites should be more host-specific to intermediate than to final hosts (Poulin 2007, Noble et al. 1989). Increased host specificity and negative fitness effects, imply that host-parasite coevolution and local adaptation may be more likely between parasites and their intermediate hosts (Lively et al. 2004).

Moreover, in host-parasite coevolution, the species with the higher dispersal rates is predicted to locally adapt to the other (Gardon and Nuismer 2009, Greischar and Koskella 2007, Morgan et al. 2005). This theoretical result is contrary to our usual expectation that dispersal and gene flow homogenize populations and counter-act divergent selection (Lenormand 2002). But in antagonistically interacting species, gene flow (within moderation) provides genetic diversity that aids in adapting to the opposing species (Gardon and Nuismer 2009). Parasite dispersal rates are usually higher than their hosts’ (Mazé-Guilmo at al. 2016, Hoeksema and Forde 2008), so parasites should be more locally adapted to their hosts than vice-versa. Moreover, hermaphroditic parasites can have higher reproductive success (i.e., it can fertilize other individual’s eggs and at the same time receive sperm to fertilize its eggs) and higher dispersion rates, both of which can lead to increased local adaptation (Mazé-Guilmo et al. 2016, Hoeksema and Forde 2008).

Combining all the propositions above (i.e., hosts-specificity, negative fitness effects on their intermediate hosts, and higher dispersal rates), parasites with complex life cycles, especially those that are hermaphroditic, should be often locally adapted to their intermediate hosts.

We tested the above prediction using the hermaphroditic tapeworm *Schistosoma solidus* (Eucestoda: Pseudophyllidea) and its first intermediate hosts, freshwater cyclopoid copepods. This tapeworm is found mainly in Holarctic lakes. It has copepods and threespine sticklebacks (*Gasterosteus aculeatus*) as first and second intermediate hosts (Barber and Scharsack 2009, Dubinina 1980) and the final hosts are warm-blooded vertebrates, usually fish-eating birds. The tapeworm reproduces sexually in the finals hosts’ intestines and its eggs are dispersed with these hosts’ feces, so the tapeworm has higher dispersal rates than its first two intermediate hosts which rarely disperse between even adjacent lakes (Caldera and Bolnick 2008). The tapeworm can be bred in-vitro, making it an excellent laboratory system for host-parasite studies (Barber 2013, Barber and Scharsack 2009, Smyth 1990). The tapeworm is not host specific to its final hosts, infecting several species of birds and even fish-eating mammals like otters (Hoberg et al. 1997, Dubinina 1980). However, the tapeworm is very host-specific to the stickleback (Barber 2013, Dubinina 1980). The tapeworm affects negatively the fitness of the fish (Weber et al. 2017b and references therein), and is locally adapted to this host (Hafer 1017, Kalbe et al. 2016). In laboratory infections, this tapeworm had negative fitness consequences to lab-reared *Macrocyclops albidus* copepods (Benesh 2010, Wedekind 1997); however, no work has been done on wild copepod species that are sympatric with the tapeworm to establish host-specificity and local adaptation, as has been done with stickleback.

We anticipate that this tapeworm would be similarly host specific and locally adapted to their copepod hosts as in their stickleback host. To test this hypothesis, we used reciprocal infection trials using factorial combinations of *S. solidus* tapeworms and native copepod species collected from lakes on Vancouver Island. We measured local adaptation through mainly infection rates in copepods by local (same lake) and foreign (different lake) tapeworms and by intensity (number of parasites inside hosts) in the infected copepods. To measure host specificity, we infected different copepod genera with the tapeworm and measured infection success in each genus. Results indicate that there was no local adaptation by the tapeworm in the copepods, but there was host specificity as a specific crustacean genus had overall higher infection rates than another used in this experiment.

## Material and methods

### Copepod colonies

We used copepods from established laboratory colonies from five lakes on Vancouver Island (Boot, Echo, Gosling, Lawier, and Roberts Lakes. The coordinates for these lakes are in supplementary Table 1). These colonies were established from plankton tows collected on September 15, 2017, and June 24, 2018. Colonies were kept in five gallon buckets at 20°C and under 16:8 hrs light:dark to simulate summer conditions in Vancouver Island until the start of the experiment on October 20 2018. We fed copepods in each bucket weekly with ∼500mL of *Paramecium caudatum* and mixed rotifer cultures plus a ground protozoan pellet, both from Carolina Biological Supply Company (Burlington, NC). We also added 10-20 autoclaved wheat seeds once a month to each bucket for bacterial growth, which contributed to the copepod and paramecium diets. Before the start of the experiment, we identified each lake’s copepods to species level under a dissecting scope and using the Image-Based Key to the Zooplankton of North America (Haney 2013).

**Table 1:** Inclusion frequencies and effect sizes of each term in the ensemble model for intensity (tapeworm count in infected copepods) and infection rate (prevalence). Effect sizes are provided as medians with 95% credible intervals and were calculated over the portion of the ensemble posterior where terms were present. Pooled terms indicate the combined effects of all copepod or tapeworm terms that were present in the model. Random effects are noted with (RE). Effect sizes for intensity and infection rate should not be compared, as they are in different units (counts and proportions, respectively).

| Model Component<sup>1</sup> | Term<sup>2</sup> | Ensemble Frequency<sup>3</sup> | Effect Size<sup>4</sup> | [95% CI]<sup>5</sup> |
| --- | --- | --- | --- | --- |
| Intensity | Copepod (pooled) | 0.849 | 0.640 | [0.421, 1.289] |
| Intensity | Copepod Genus | 0.849 | 0.621 | [0.019, 2.201] |
| Intensity | Worm (pooled) | 0.817 | 0.187 | [0.027, 1.297] |
| Intensity | Worm Lake | 0.817 | 0.182 | [0.023, 1.374] |
| Intensity | Copepod Lake | 0.522 | 0.503 | [0.190, 2.464] |
| Intensity | Plate (RE) | 0.500 | 0.095 | [0.004, 0.421] |
| Intensity | Native | 0.473 | 0.126 | [0.005, 0.798] |
| Intensity | Worm Family (RE) | 0.420 | 0.113 | [0.006, 0.575] |
| Intensity | Genus x Worm Lake Interaction | 0.354 | 0.616 | [0.052, 4.309] |
| Intensity | Copepod x Worm Lake Interaction | 0.208 | 0.721 | [0.079, 7.482] |
| Infection Rate | Copepod (pooled) | 0.871 | 0.255 | [0.213, 0.364] |
| Infection Rate | Copepod Genus | 0.871 | 0.274 | [0.043, 0.385] |
| Infection Rate | Worm (pooled) | 0.840 | 0.058 | [0.020, 0.118] |
| Infection Rate | Worm Lake | 0.840 | 0.060 | [0.018, 0.128] |
| Infection Rate | Copepod Lake | 0.593 | 0.143 | [0.072, 0.242] |
| Infection Rate | Plate (RE) | 0.500 | 0.032 | [0.002, 0.078] |
| Infection Rate | Worm Family (RE) | 0.409 | 0.018 | [0.001, 0.060] |
| Infection Rate | Native | 0.372 | 0.039 | [0.002, 0.142] |
| Infection Rate | Copepod x Worm Lake Interaction | 0.235 | 0.095 | [0.045, 0.164] |
| Infection Rate | Genus x Worm Lake Interaction | 0.218 | 0.053 | [0.009, 0.175] |

<sup>1</sup> For Model Component; Infection Rate is the Prevalence of infection in copepods by the tapeworm. Intensity is the number of tapeworms inside infected copepods.
<sup>2</sup>Factor indicates the model term; Copepod (pooled) and Worm (pooled) refer to the combination of whatever copepod/worm related terms were present in each model.
<sup>3</sup>Frequency the proportion of the model ensemble that the term appears in.
<sup>4</sup>Effect size of each model term accounting for the data.
<sup>5</sup>CI: 95% credible interval of the effect size.

The laboratory colonies for each lake only had one surviving copepod species just before the start of the experiment. These were *Macrocyclops albidus* for Boot and Lawier Lakes, *Macrocyclops fuscus* for Roberts Lake, *Acanthocyclops robustus* for Echo Lake, and *Acanthocyclops brevispinosus* for Gosling Lake. All these copepods were from the order Cyclopoida.

### Tapeworm colonies

We used tapeworm eggs from three lakes in Vancouver Island (Boot, Echo, and Gosling Lakes). Lawier and Roberts Lakes lack infected stickleback fish, so tapeworms were unavailable from these two lakes; thus, we infected copepods from five lakes with tapeworms from three. The advantage of this design is that copepods from Lawier and Roberts lakes could be highly susceptible to the tapeworm due to their lesser exposure to the parasites; thus, serving as positive controls. The tapeworm eggs were collected from laboratory crosses of randomly chosen wild tapeworms obtained from infected fish, following established methods (Weber et al. 2017b, Smyth 1990). We hoped that these randomly chosen tapeworms would reflect the tapeworm genetic diversity in each lake. These crosses were done in June – Sept. 2018, and the eggs were kept at 4°C until the experiment.

### Experimental set-up

To test for tapeworm local adaptation and host specificity to copepods, we carried out a reciprocal infection experiment by exposing the copepods from each lake to local and foreign tapeworm larvae (coracidia) from three lakes (i.e., Boot, Echo, and Gosling lakes). We hatched tapeworm eggs and exposed the coracidia to copepods following published methods (Weber et al. 2017b, Smyth 1990). We used six-well plates, each well holding a different combination of copepods (n=10 individuals per well) from a lake and tapeworms (n=20 coracidia per well) from the same or a different lake (Figure 1). We used a combination of 1:2 copepod to tapeworm ratio to account for the short lifespan (∼24hr) of the parasite (Dubinina 1980). We used three tapeworm families or strains per lake. We also had six to eight wells per lake with copepods unexposed to tapeworms as negative controls to measure tapeworm exposure and infection effects on host mortality (supplementary table 2). The plates were kept in the same conditions as the copepod colonies (i.e. 20°C and 16:8hrs light:dark). We randomized the positions of the copepod-tapeworm combinations within plates, and plate locations within the incubator. We dissected each surviving copepod to ascertain infection status 17-22 days post exposure when tapeworms reached maximum size inside copepods (Dubinina 1980).

**Figure 1:**
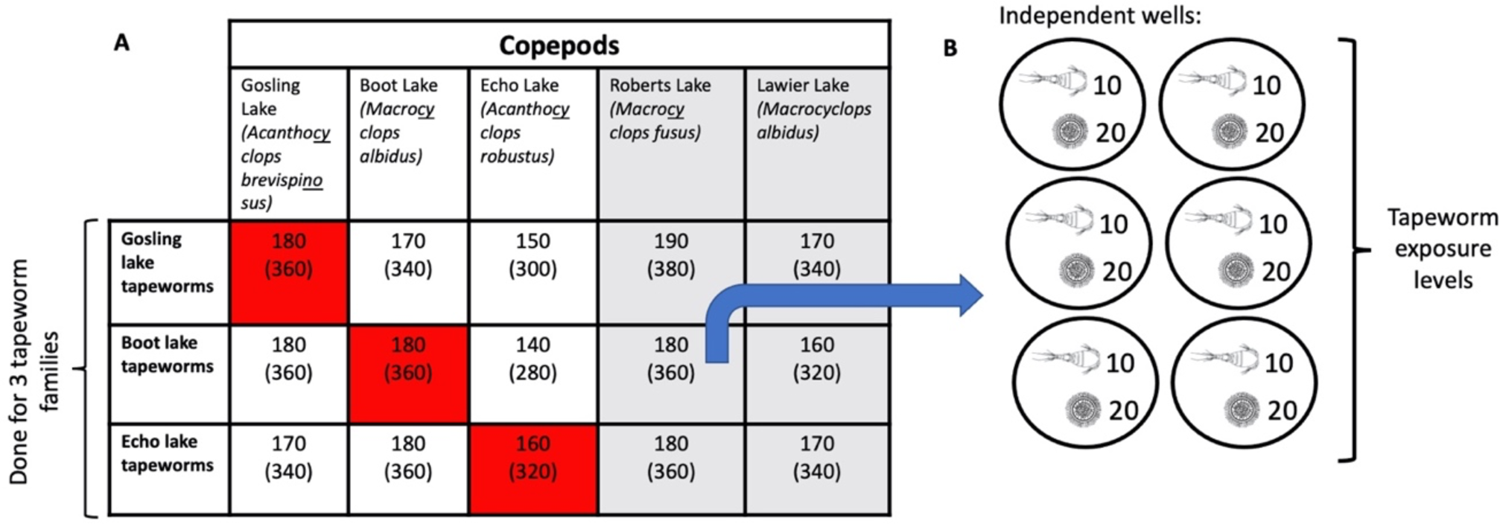
Graphical representation of the experiment setup. **A)** The combinations of the tapeworm *Schistocephalus solidus* by copepod exposures, using three tapeworm families per lake; red squares indicate tapeworms exposed to sympatric copepods. Roberts and Lawier Lakes are shaded in grey representing control lakes where the tapeworm is lacking in threespine sticklebacks. The numbers inside each square represent total numbers of copepods and tapeworms used (the latter in parenthesis). Names of the copepod species used are below each lake’s names. **B)** A diagram of how each tapeworm family was exposed to each lake’s copepods (in this example Boot Lake tapeworms to Roberts Lake copepods): in six different wells from different 6-well plates, each with 10 copepods exposed to 20 tapeworm larvae. All well positions for all exposures in panel A were randomized in the 6-well plates, and the position for each 6-well plates were also randomized in the experimental room.

In total, we used 49 6-well plates, exposing 2,890 copepods (10 per well) with 5,780 tapeworms (20 per well, nine families in total, three per lake. See supplementary table 2). At the end of experiment, 1622 exposed and 330 control copepods survived. Exposure to tapeworms did not affect copepod survival (P value = 0.996, supplementary figure 2). The survival rate for copepods in the experiment was 56%.

**Figure 2.**
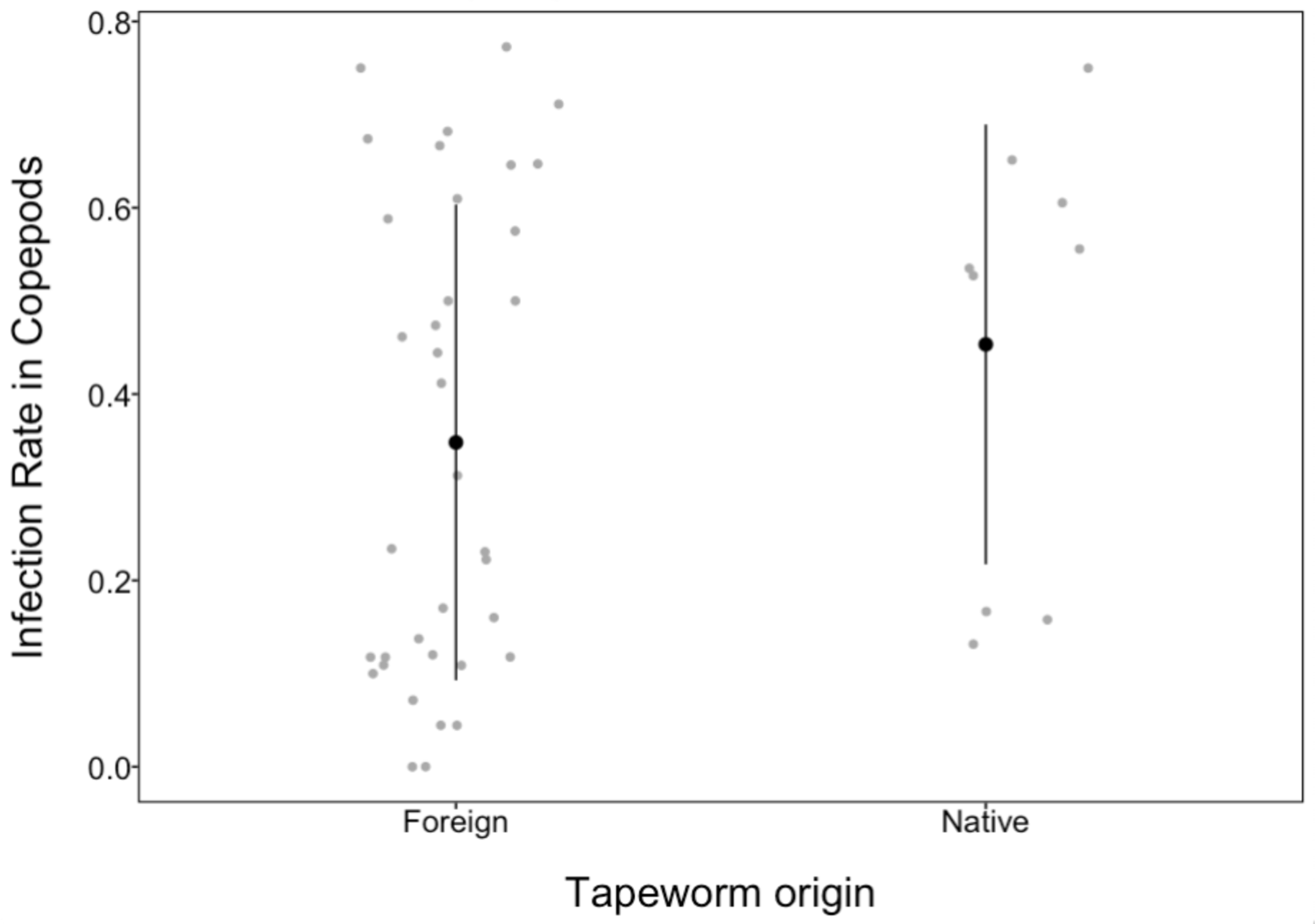
Overall, infection rates on copepods by local or native tapeworms (i.e. where the *S. solidus* tapeworms are from the same lakes as the copepods) are very similar to that of foreign tapeworms.

### Bayesian analysis (data analysis)

We used mixed-effect hurdle models to simultaneously estimate the effect of copepod and parasite origin on infection rate (prevalence) and intensity (number of worms per successfully infected copepod). Conceptually, these models combine a logistic regression on parasite presence/absence with a truncated Poisson regression on non-zero parasite counts. Our models considered tapeworm lake and its interaction with either copepod genus or lake as fixed effects; we also included an indicator for whether the tapeworm and copepod were from the same lake (i.e. “native”). Plate number and tapeworm lake were included as random effects. The full model contains all of these terms as predictors for both prevalence and incidence. We created a series of reduced models from a list of all possible combinations of predictors, excluding models that contained interactions without their main effects, copepod lake without genus, and tapeworm family without lake.

We fit all models with the *brms* package in R v. 4.0.4 (Bürkner 2018, R Core Team 2018). The predictive value of each model was determined with Bayesian stacking weights calculated by the *loo* package (Yao et al. 2018); conceptually, this is similar to AIC model weighting. A combined ensemble was created by pooling a weighted sample of each model’s posterior distribution. We defined effect sizes as the standard deviation of a term’s marginal effects at each posterior sample from a model where the term was included. Prior specification and other details are provided in the supplementary material section.

## Results

In this paper we tested if the tapeworm *S. solidus* is locally adapted and host specific to their copepod hosts. My results indicate that the tapeworm is not locally adapted to the copepods (figure 2), but it might be more host specific to a genus of copepods as rates of infection and intensity (number of parasites inside infected hosts) were higher in *Acanthocyclops* than *Macrocyclops* copepods (figures 3 to 5).

**Figure 3:**
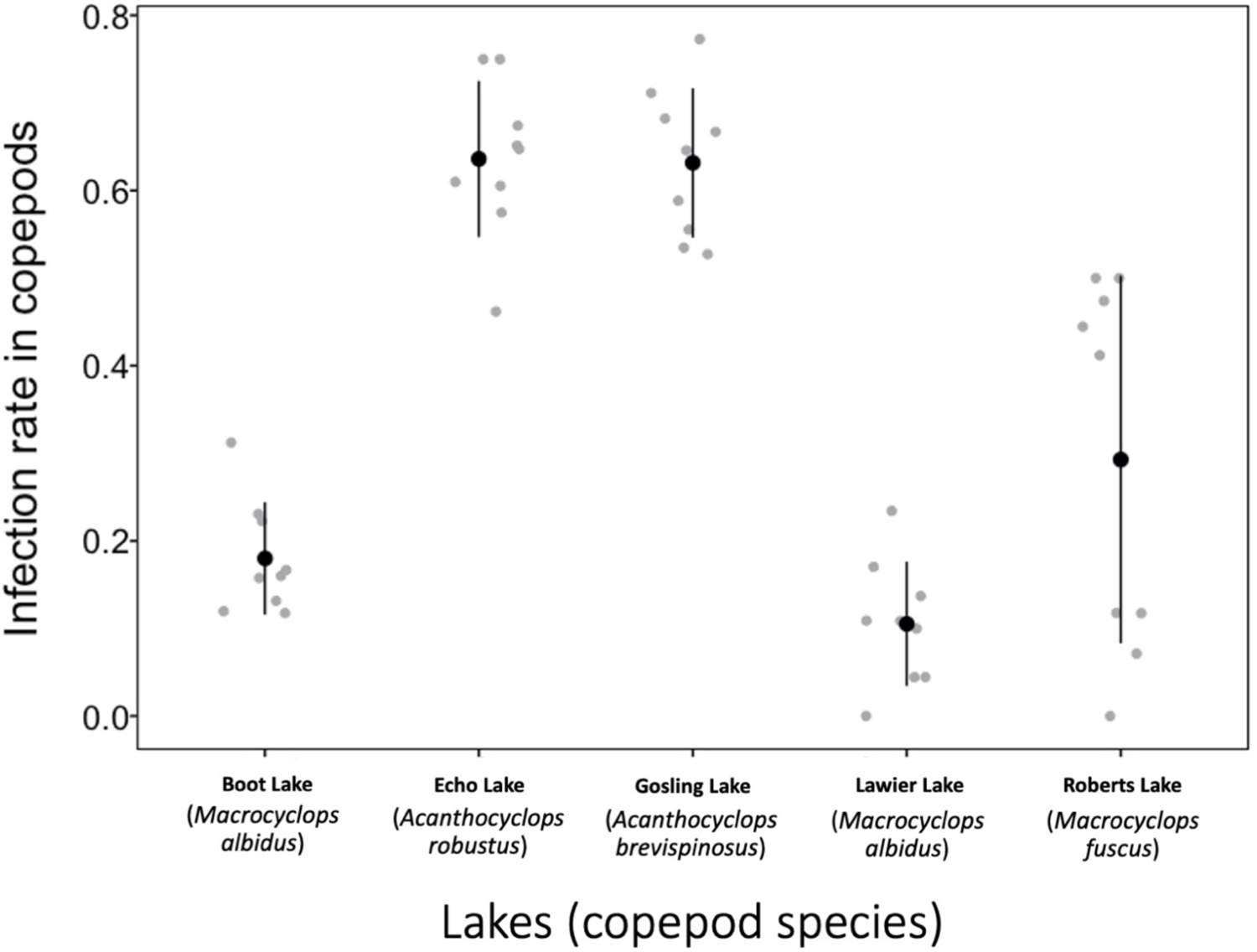
Infection results indicate that copepods from Echo and Gosling Lakes were three to six times more susceptible to infection by the tapeworm than those of the other three lakes. The scientific names of the copepods from each lake are in parenthesis under the lake names. The tapeworm is not found in Lawier and Roberts lakes (at least from stickleback fish surveys).

**Figure 4.**
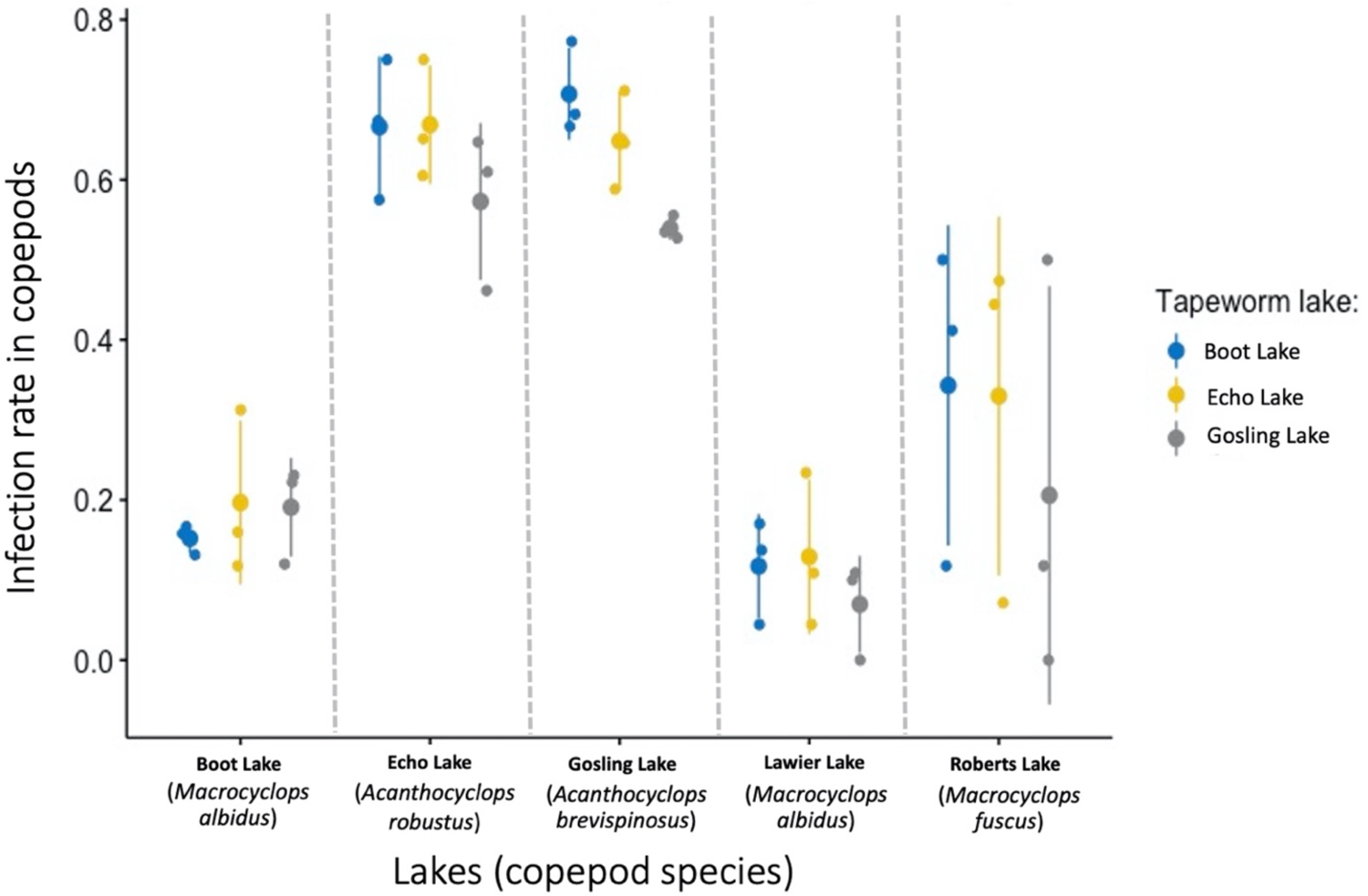
The infection rates in copepods were similar among all the tapeworm families or strains from the three lakes used. Again, copepods from Echo and Gosling Lakes were three to six times more susceptible to infection than those of the other three lakes (see figure 2). The scientific names of the copepods from each lake are in parenthesis under the lake names. The tapeworm is not found in Lawier and Roberts lakes (at least from stickleback fish surveys).

**Figure 5.**
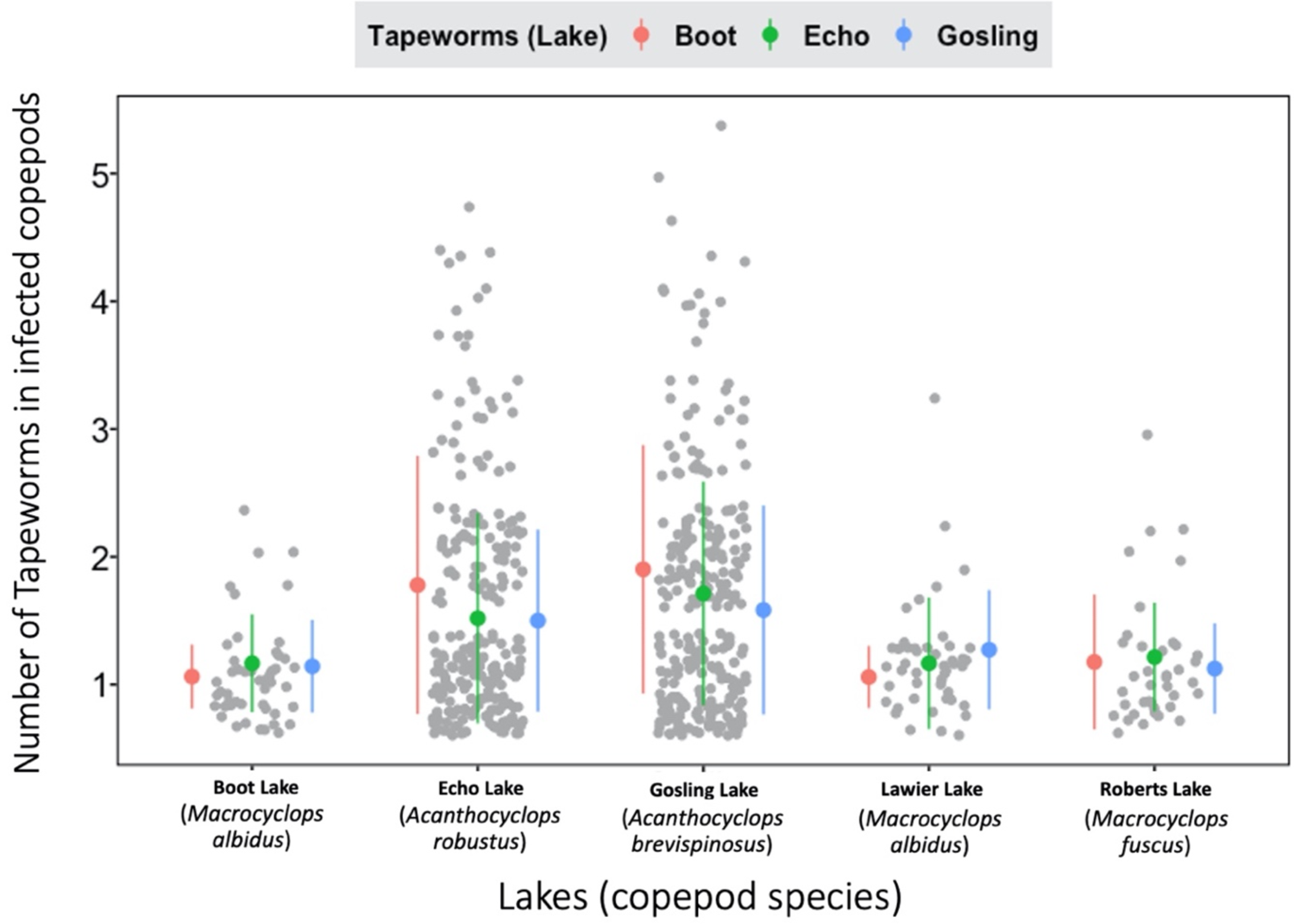
The infected copepods from Echo and Gosling Lakes also had slightly more parasites on average than the ones from the other three lakes. Again, the averages were very similar in the three tapeworm strains from the three lakes used. The scientific names of the copepods from each lake are in parenthesis under the lake names. The tapeworm is not found in Lawier and Roberts lakes (at least from stickleback fish surveys).

### Results from the Bayesian analysis

The ensemble mixed-effect hurdle model contained 1.37 million posterior samples, with 4,450 different models contributing at least one sample. No single model had a stacking weight higher than 0.3%; however, both copepod and tapeworm origins contributed to over 80% of both the intensity and infection rate model components (Table 1). For both model components, copepods had the largest effect size of any term (intensity: 0.640 [0.421, 1.289]; infection rate: 0.255 [0.213, 0.364]; brackets signify 95% credible interval). Copepod effects can be decomposed into genus and lake of origin, with lake nested in genus; 61% of posterior samples with copepod genus terms also contained copepod lake. Copepod genus effect sizes were generally smaller when lake effects were also present (intensity effect sizes: 0.712 [0.518, 0.958] without lake, 0.342 [0.011, 2.714] with lake; infection rate: 0.338 [0.303, 0.375] without lake, 0.210 [0.031, 0.395] with lake; supplementary figure 1).

Copepod by tapeworm interactions (the typical test for local adaptation) had the lowest ensemble inclusion frequencies for both the intensity and infection rate model components (Table 1), and their effect sizes when present had wide, noisy posterior distributions. The ‘native’ effect (indicating copepods and tapeworms from the same lake) had higher inclusion but consistently small effect sizes; we interpreted this as insufficient evidence for local adaptation. All of these effects had lower inclusion rates than the 6-well plates used for in the experiment.

We also ran mix-effect linear and GLM models in R (R Core Team) to supplement the analyses and results above. For these analyses, the best predictors for infection rate were the copepod and tapeworm lakes, and the best predictors for intensity in infected copepods was copepod lake. These results did not differ considerably from the best Bayesian mixed-effect hurdle models above, suggesting our results are robust to either choice of analytical method. For more details on the mix-effect and GLM models and results, see supplementary material.

As mentioned in the Bayesian results section, there was not enough evidence for local adaptation of the tapeworm to their copepod hosts. This can also be seen in figure 2, where infection rates by the tapeworm on local (from the same lake) and foreign (from different lakes) copepods were very similar. However, there was evidence of host specificity as copepod genus was a strong predictor in infection rate and infection intensity in the crustacean. For example, copepods from Echo and Gosling lakes (both of the genus *Acanthocyclops*) were three to six times more susceptible to infection than the other copepod genus (*Marcocyclops*) from the three remaining lakes (figure 3). This was true for all tapeworm strains used (figure 4). Moreover, the infected copepods from Echo and Gosling lakes (again both of the genus *Acanthocyclops*) also had between 0.3 to 0.5 times more tapeworms than those (of the genus *Marcocyclops)* from the other three lakes (figure 5). This accounts for the relatively high effect sizes of the copepod genus factor in the Bayesian analysis.

## Discussion

We tested for local adaptation and host specificity of the tapeworm *S. solidus* from three lakes in Vancouver Island to copepods from the same plus two more lakes where the tapeworm is absent (Weber et al. 2017b, personal observations). Researchers argue that parasites with complex life cycles should be more host-specific (Poulin 2007, Nobel et al. 1989), and that parasites with higher dispersal rates should locally adapt to their hosts (Barber and Scharsack 2009, Morgan et al. 2009). Thus, the *S. solidus* tapeworm, being a parasite with a complex life cycle and having higher dispersal rates than their intermediate hosts (Dubinina 1980), should show local adaptation and host specificity to its copepod hosts in a similar fashion to the tapeworm’s second intermediate host (i.e. threespine sticklebacks [Hafer 2018, Weber et al. 2017a, Kalbe et al. 2016]).

However, our results indicate that there was no evidence of differences between infection rates by local and foreign tapeworms on the copepods (figure 2). Our experiment also shows that copepods from Echo and Gosling Lakes (genus *Acanthocyclops*) were more susceptible to *S. solidus* tapeworm infection than the ones from the other three lakes (genera *Macrocyclops*) (figure 3), and these copepods also had slightly more tapeworms when infected (figure 5). These infection and intensity rates were very similar among the different tapeworm strains from the three lakes used (figures 4 and 5). Thus, at least for this parasite-host system, we did not observe local adaptation by the tapeworm to the copepods.

Instead, the success of the tapeworm within a given lake depended mostly on whether a copepod genus (*Acanthocyclops*) was present. Variation in zooplankton community structure, between lakes, means that tapeworms will be locally maladapted to lakes with *Macrocyclops* spp copepods. The higher susceptibility of Echo and Gosling Lakes’ copepods (of the genus *Acanthocyclops*) to the tapeworm explains why copepod and tapeworm lake variables in our models fit most the data.

To emphasize more the lack of local adaptation in our experiments, Boot and Lawier Lakes had the same species of copepods (*Macrocyclops albidus*), but both lakes’ copepods had very similar infections rates by the three strains of tapeworms used (figure 4). Specifically, Boot Lake tapeworms are no more (or less) effective at infecting Boot Lake *M. albidus* than they are at infected Lawier Lake *M. albidus* (a home-versus-away criterion for local (mal)adaptation). Nor are the Boot Lake tapeworms any better (or worse) at infecting their native Boot Lake copepods, relative to tapeworms from two other lakes (a native versus immigrant criterion for local (mal)adaptation). Thus, for both lakes, the infection rate by local tapeworms was not significantly different to that of foreign tapeworms.

Local adaptation aside, our experiments show the tapeworm is clearly capable of infecting multiple copepod genera, but it is most efficient at infecting a particular genus. The copepods with the highest infection rates (those from Echo and Gosling lakes) were from the same genus (i.e. *Acanthocyclops*). This was true regardless of whether the tapeworms were taken from a lake dominated by *Acanthocyclops*, or not. Currently, *M. albidus* copepods are used for experimental infections in sticklebacks (Weber et al. 2017a and 2017b, Barber 2013, Benesh 2010, Wedekind 1997, Smyth 1990); perhaps, future work should employ *Acanthocyclop* species instead to maximize resources and time for better results. Weber et al. (Weber et al. 2017a) argued that to understand the patchiness of the tapeworm infections in stickleback populations, more data is needed on ecological processes like parasite encounter rates and abundance of suitable primary hosts (copepods). Although, the primary reason for the different stickleback infection levels in the lakes sampled was due to recent evolution of the fish’s immunology (Weber et al. 2017b), copepod infectivity might still play a role. We sampled in the same lakes for this work, so we can comment on the stickleback infection levels to our copepod infection levels. The high copepod infection levels in Gosling and Echo Lakes might contribute to the high stickleback infection levels in these lakes (figs 1 in Weber et al. 2017a and 2017b). And even if wild sticklebacks in Roberts Lake lack tapeworms, our experiments here show that this lake’s copepods can get infected, validating the hypothesis that this lake’s fish are exposed to tapeworms in the wild but still have zero infections due to recently evolved immunological mechanisms to combat infections (Weber et al. 2017b).

## Supplementary material

**Supplementary table 1:** copepod collection dates and locality:

| Lake | Longitude | Latitude | Collection date |
| --- | --- | --- | --- |
| Boot Lake | 50.05503 | -125.526 | 9/16/17, 6/24/18 |
| Echo Lake | 49.98765 | -125.411 | 9/15/17, 6/24/18 |
| Lawier Lake | 50.083144 | -125.515052 | 9/15/17, 6/24/18 |
| Roberts Lake | 50.216390 | -125.544687 | 9/15/17, 6/24/18 |
| Gosling Lake | 50.04592 | -125.501 | 9/16/17, 6/24/18 |

**Supplementary table 2:** Tapeworm families used for the experiments and number of exposures (i.e. wells in a 6-well plate) for each lake’s copepods (reminder: each well had 10 copepods exposed to 20 tapeworms):

| Tapeworm families<sup>1</sup> | Gosling Lake copepods | Roberts Lake copepods | Lawier Lake copepods | Echo Lake copepods | Boot Lake copepods |
| --- | --- | --- | --- | --- | --- |
| Control | 6 | 6 | 6 | 8 | 7 |
| Boot 11Bx1A (6/17/18) | 6 | 6 | 6 | 5 | 6 |
| Boot 2Ax2C (6/17/18) | 6 | 6 | 5 | 4 | 6 |
| Boot bulk (7/3/18) | 6 | 6 | 5 | 5 | 6 |
| Echo bulk (6/15/18) | 6 | 6 | 6 | 5 | 6 |
| Echo 3Ax1A (6/10/18) | 6 | 6 | 6 | 5 | 6 |
| Echo 27Ax31A (6/17/18) | 5 | 6 | 5 | 6 | 6 |
| Gosling 7Ax1A (6/10/18) | 6 | 6 | 6 | 5 | 6 |
| Gosling 10Ax12A (6/17/18) | 6 | 7 | 5 | 5 | 5 |
| Gosling 2 (9/22/18) | 6 | 6 | 6 | 5 | 6 |

<sup>1</sup> Tapeworm families used in the experiment; 3 families per lake; dates in the parentheses below each family indicate the time the tapeworm eggs were harvested in the lab for the experiment. Control indicates no tapeworms were used to exposed the copepods (i.e. negative control); this is to evaluate the survivorship of the copepods in the wells during the experiment.

**Supplementary figure 1:**
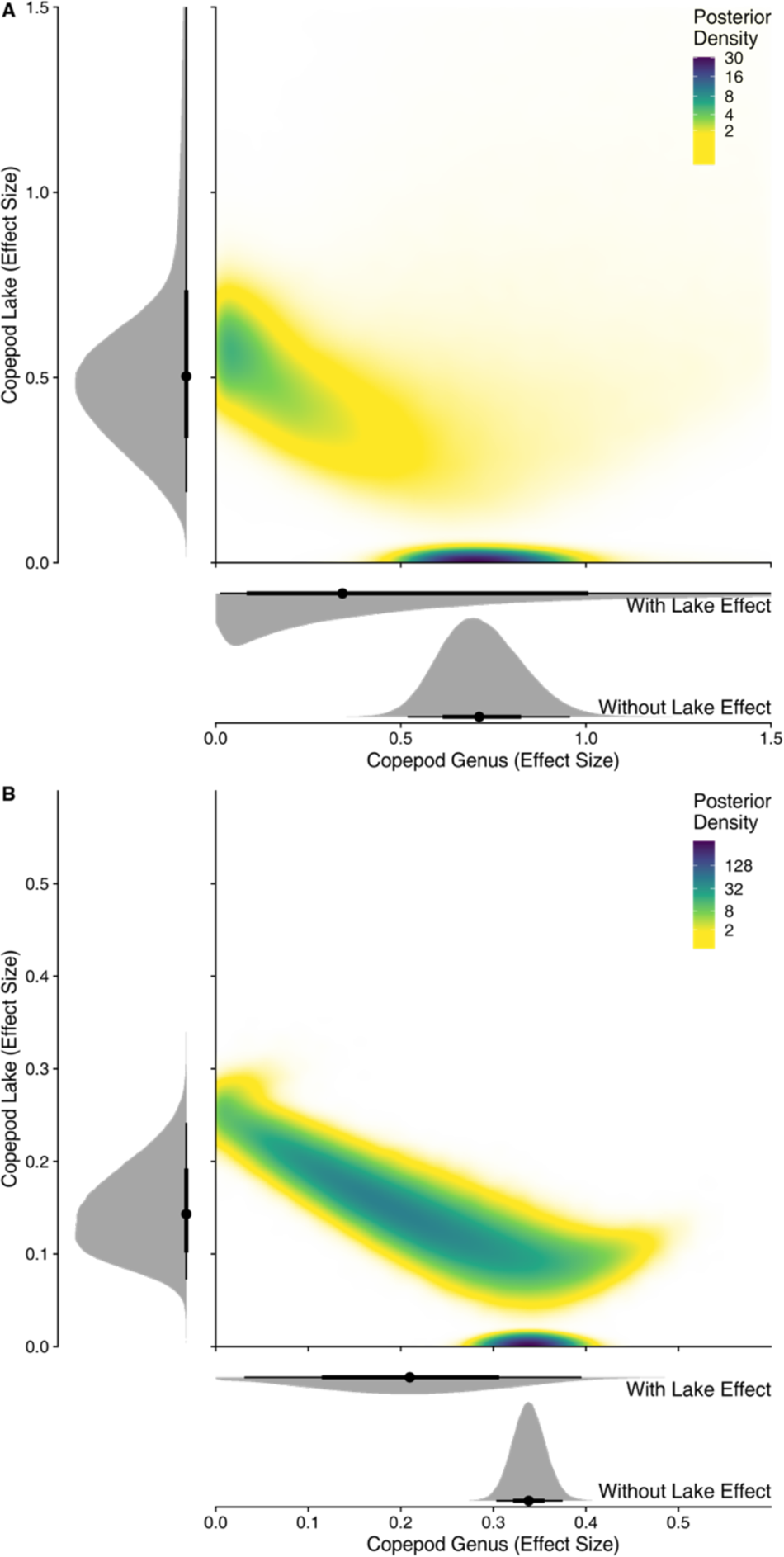
Distribution of effect sizes for Copepod Genus and Copepod Lake for the infection intensity. (**A**) and rate (**B**) model components. Both panels include the marginal effects of copepod lake (left), the marginal effects of copepod genus conditioned on whether lake was included in the model (bottom), and their bivariate distributions (upper right). For both terms, the genus effect increases when the lake effect declines or is absent; this is particularly notable for the intensity component.

**Supplementary figure 2:**
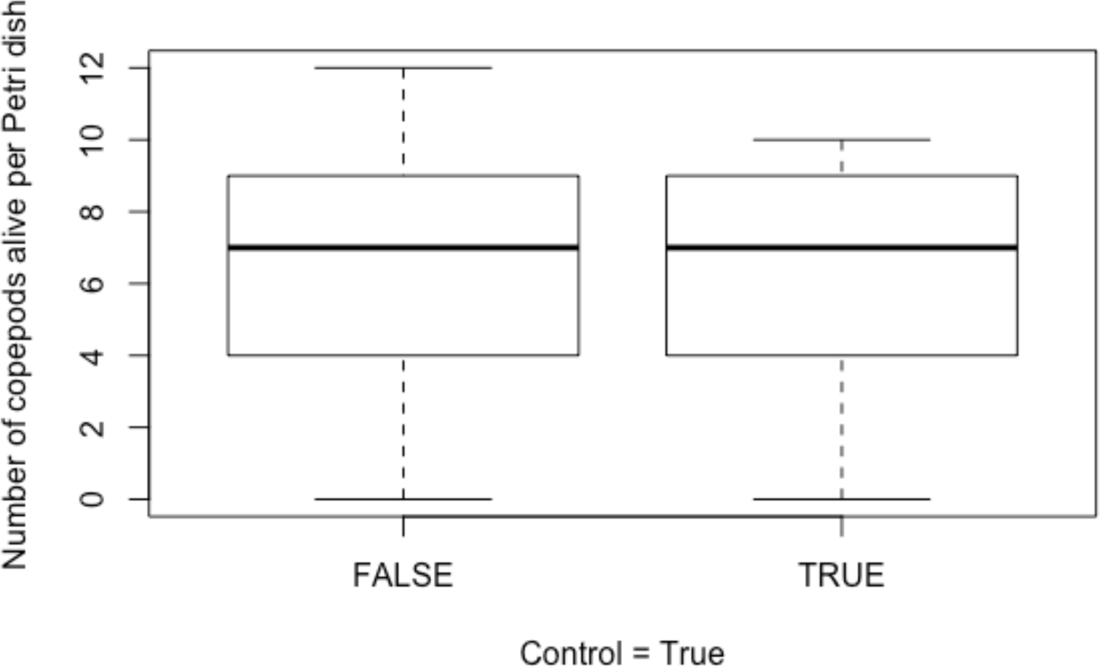
The number of copepods alive after termination of experiment did not differ significantly between those exposed to the tapeworm and those that were not (i.e., control) [P value = 0.996, see more details of analysis in Supplementary mix-effect linear and GLM model analyses below]

### Priors and iterations used in the mixed-effect hurdle analysis

Each model was run for 4 chains with 1000 warmup and 1000 sampling iterations each. We checked model convergence by verifying N_eff > 1000, R-hat > 1.01, and all Hamiltonian Monte-Carlo diagnostics were acceptable.

Our prior distributions were Normal(mean = 0, sd = 6) priors for the intercepts of both model components. For incidence, we used Normal(0,1) priors for fixed effect coefficients and half-t(df = 7, mean = 0, scale = 1) priors for the standard deviation of the random effects. For the prevalence model, our priors were Normal(0, 1.5) for fixed effects and half-t(7, 0, 1.5) for the random effects. These priors were selected because they were flexible enough to allow for large effects but conservative enough to avoid spurious results.

Note: The complete R script for the mixed-effect hurdle analyses is in Christopher Peterson’s GitHub (https://github.com/Christopher-Peterson/copepod_worm_adapt).

### Supplementary mix-effect linear and GLM model analyses

I) Analyzing if there was any difference on survival rate between copepods exposed to tapeworms and the not-exposed ones (i.e. control):

> model15 = glm(cop.alive ∼ is_control, data = control_df)

> anova(model15, test = “LRT”)

Analysis of Deviance Table

Model: Poisson, link: log

Response: cop.alive

Terms added sequentially (first to last)

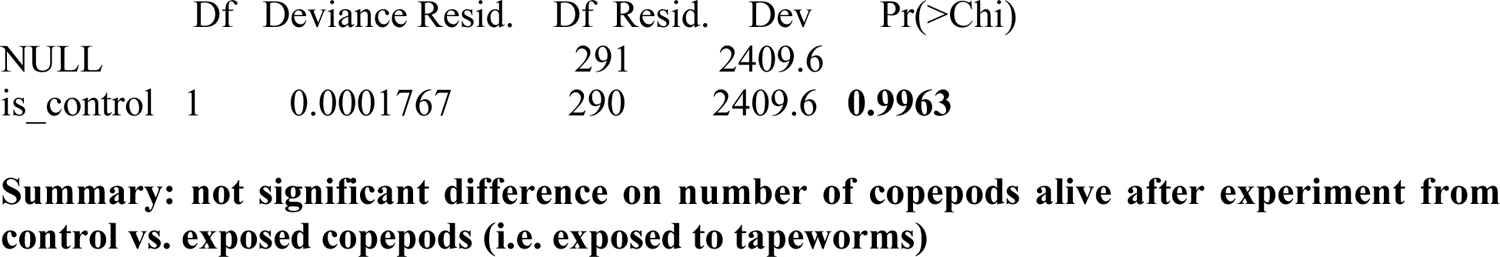

> model14 = glm(cop.death.numb ∼ is_control, data = control_df, family = poisson())

> anova(model14, test = “LRT”)

Analysis of Deviance Table

Model: poisson, link: log

Response: cop.death.numb

Terms added sequentially (first to last)

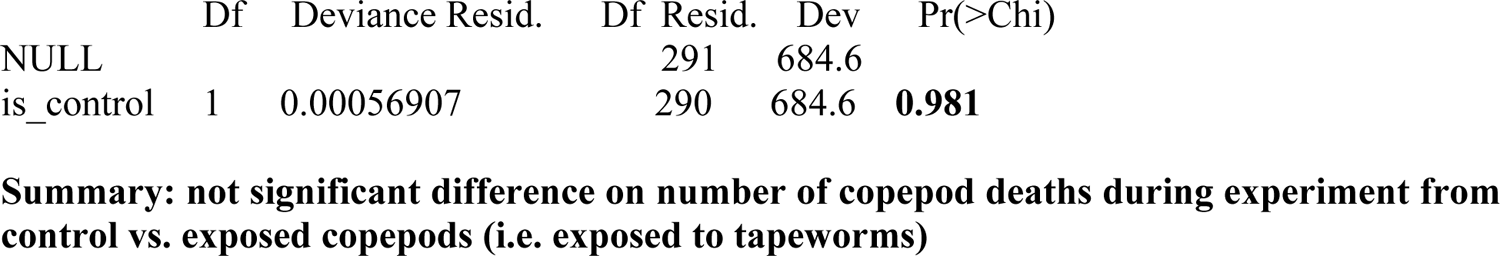

**II). GLM and GLMM analyses companion to the Bayesian analysis**

Using lme4 version 1.1-13 package for R

> summary(model1)

Generalized linear mixed model fit by maximum likelihood (Laplace Approximation) [glmerMod]

Family: binomial (logit)

Model 1: infected.yes.no ∼ cop.lake * worm.lake + (1 | plate)

Data: copepods

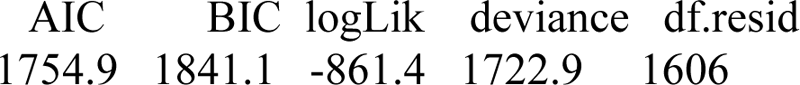

Scaled residuals:

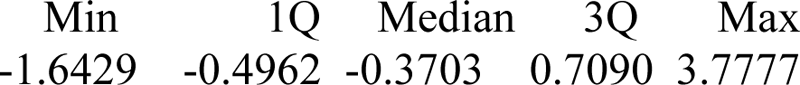

Random effects:

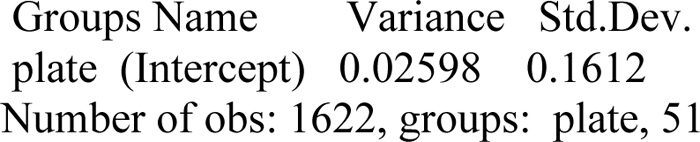

Fixed effects:

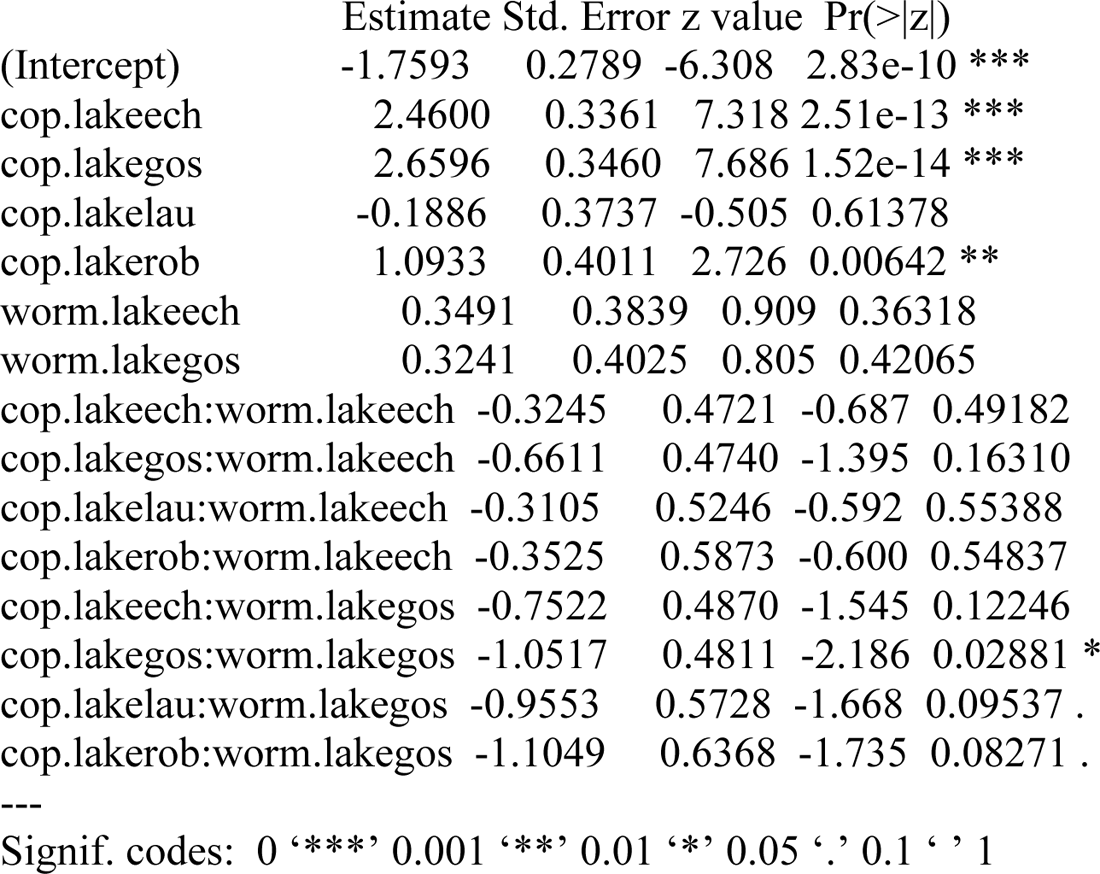

Correlation matrix not shown by default, as p = 15 > 12.

Use print(x, correlation=TRUE) or vcov(x) if you need it

convergence code: 0

Model failed to converge with max|grad| = 0.00320527 (tol = 0.001, component 1)

> anova(model1,test=“LRT”)

Analysis of Variance Table

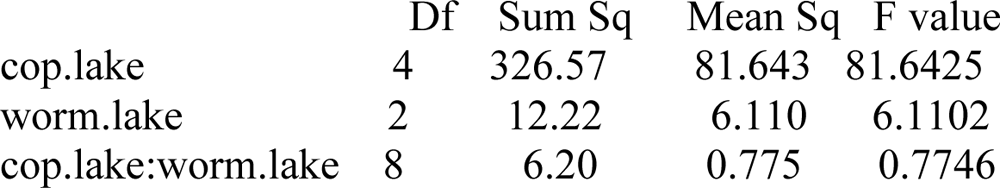

**2) Doing another model without the interaction of copepod lakes and worm lakes:**

> model2 <-glmer(infected.yes.no ∼ cop.lake + worm.lake + (1|plate),

+ data=copepods, family=“binomial”) #model 2 does not have an interaction

> #comparing models 1 and 2:

> anova(model1, model2)

Data: copepods

Models:

model2: infected.yes.no ∼ cop.lake + worm.lake + (1 | plate)

model1: infected.yes.no ∼ cop.lake * worm.lake + (1 | plate)

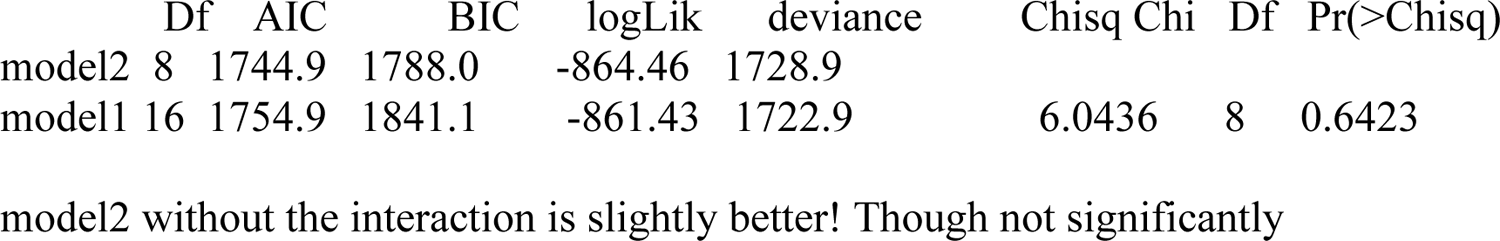

**3) model 6, testing for GLM without taking into account the random variable (no random effect)**

> model6 <-glm(infected.yes.no ∼ cop.lake + worm.lake + cop.lake*worm.lake, + data = copepods,family=“binomial”)

> summary(model6)

Call:

glm(formula = infected.yes.no ∼ cop.lake + worm.lake + cop.lake * worm.lake, family = “binomial”, data = copepods)

Deviance Residuals:

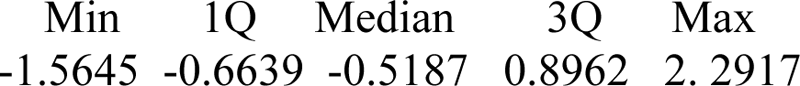

Coefficients:

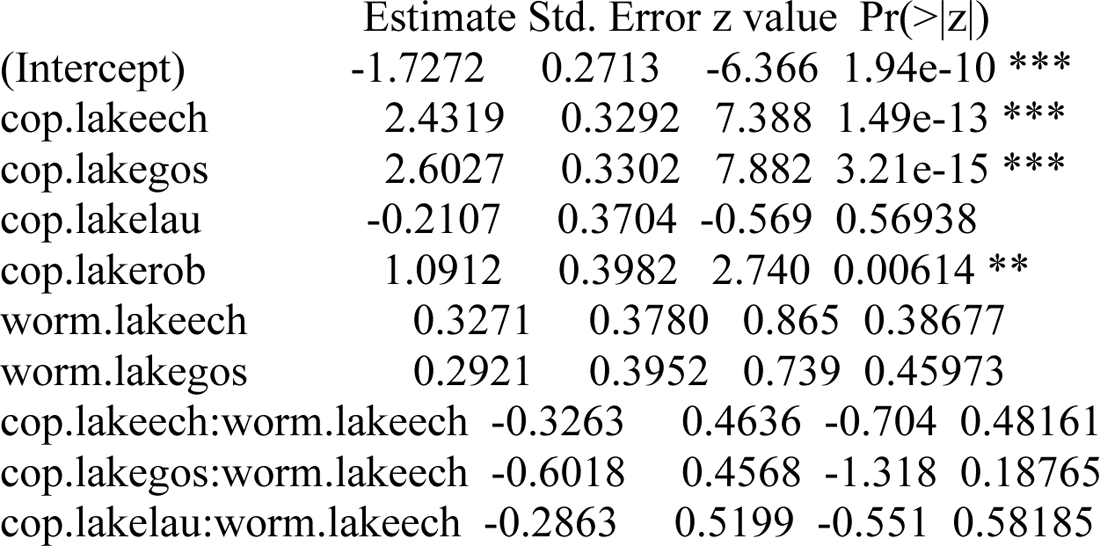
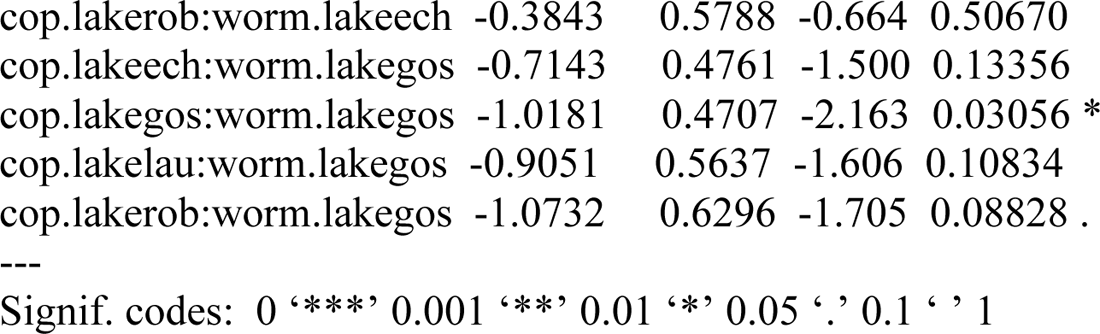

(Dispersion parameter for binomial family taken to be 1)

Null deviance: 2167.1 on 1621 degrees of freedom

Residual deviance: 1723.3 on 1607 degrees of freedom

AIC: 1753.3

Number of Fisher Scoring iterations: 5

> anova(model6,test=“LRT”) #“LRT” is to get P-values here to check quickly for significance for the variables in the model

Analysis of Deviance Table

Model: binomial, link: logit

Response: infected.yes.no

Terms added sequentially (first to last)

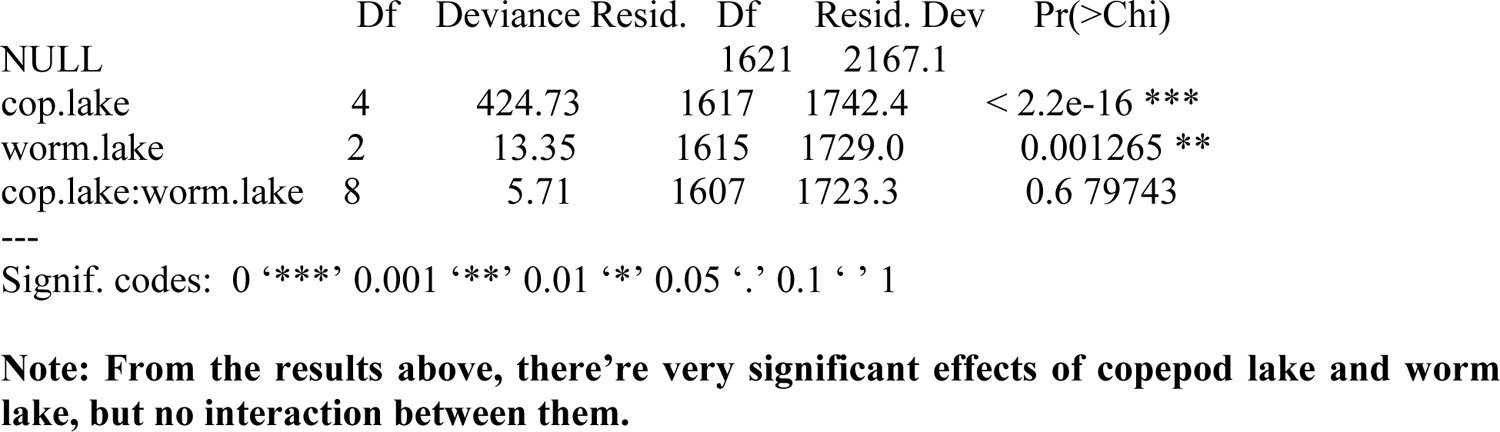

**4) testing for effect of plates: (summary answer after running the model below: plate does not matter)**

> model7 <-glm(infected.yes.no ∼ cop.lake + worm.lake + cop.lake*worm.lake + plate,

+ data=copepods,family=“binomial”)

> anova(model6, model7) #does not give AIC comparisons, so this kind of useless Analysis of Deviance Table

Model 1: infected.yes.no ∼ cop.lake + worm.lake + cop.lake * worm.lake

Model 2: infected.yes.no ∼ cop.lake + worm.lake + cop.lake * worm.lake + plate

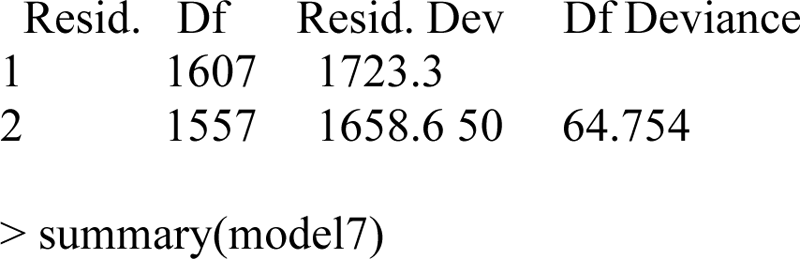

Call:

glm(formula = infected.yes.no ∼ cop.lake + worm.lake + cop.lake * worm.lake + plate, family = “binomial”, data = copepods)

Deviance Residuals:

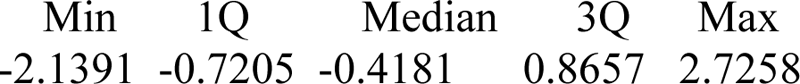

Coefficients:

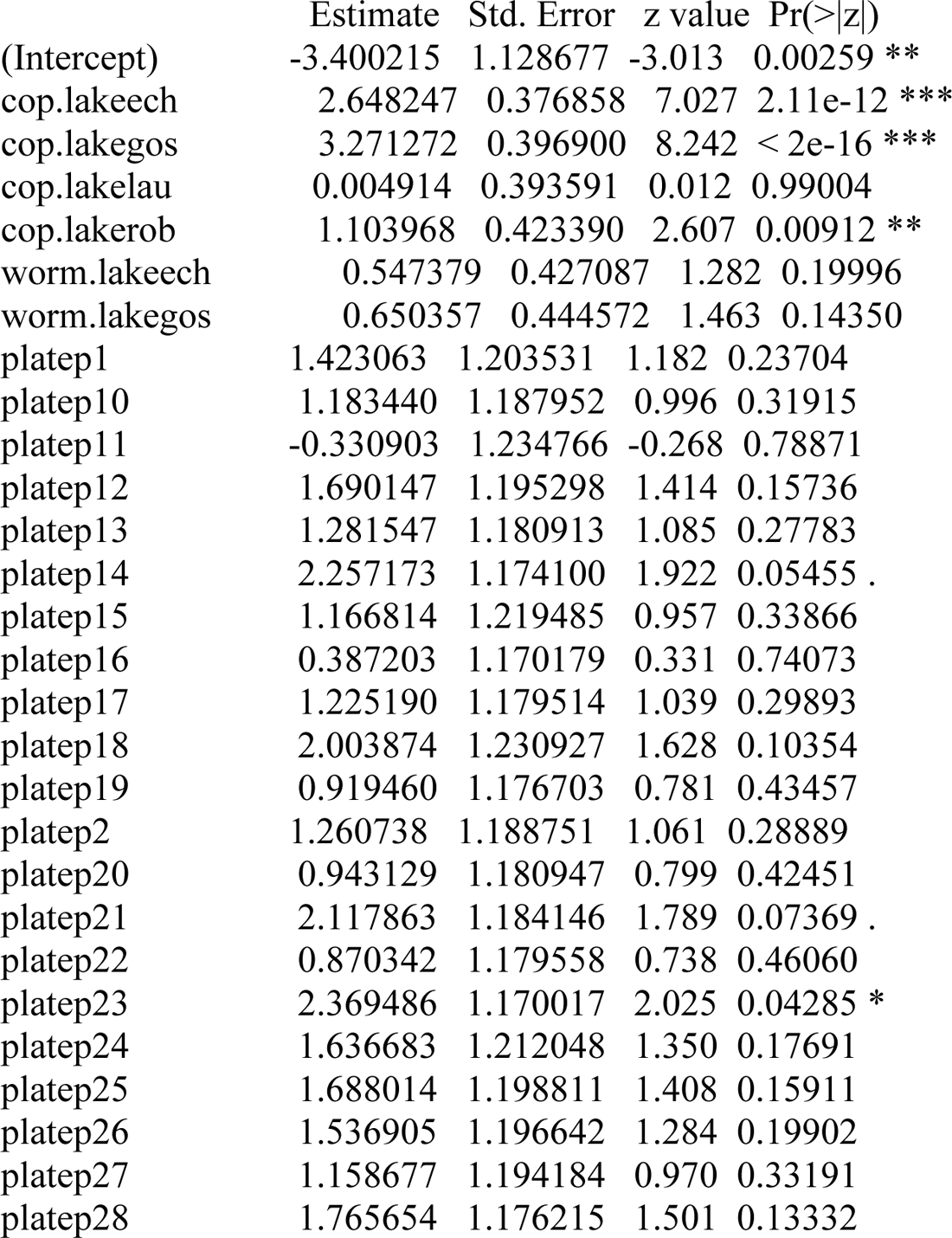
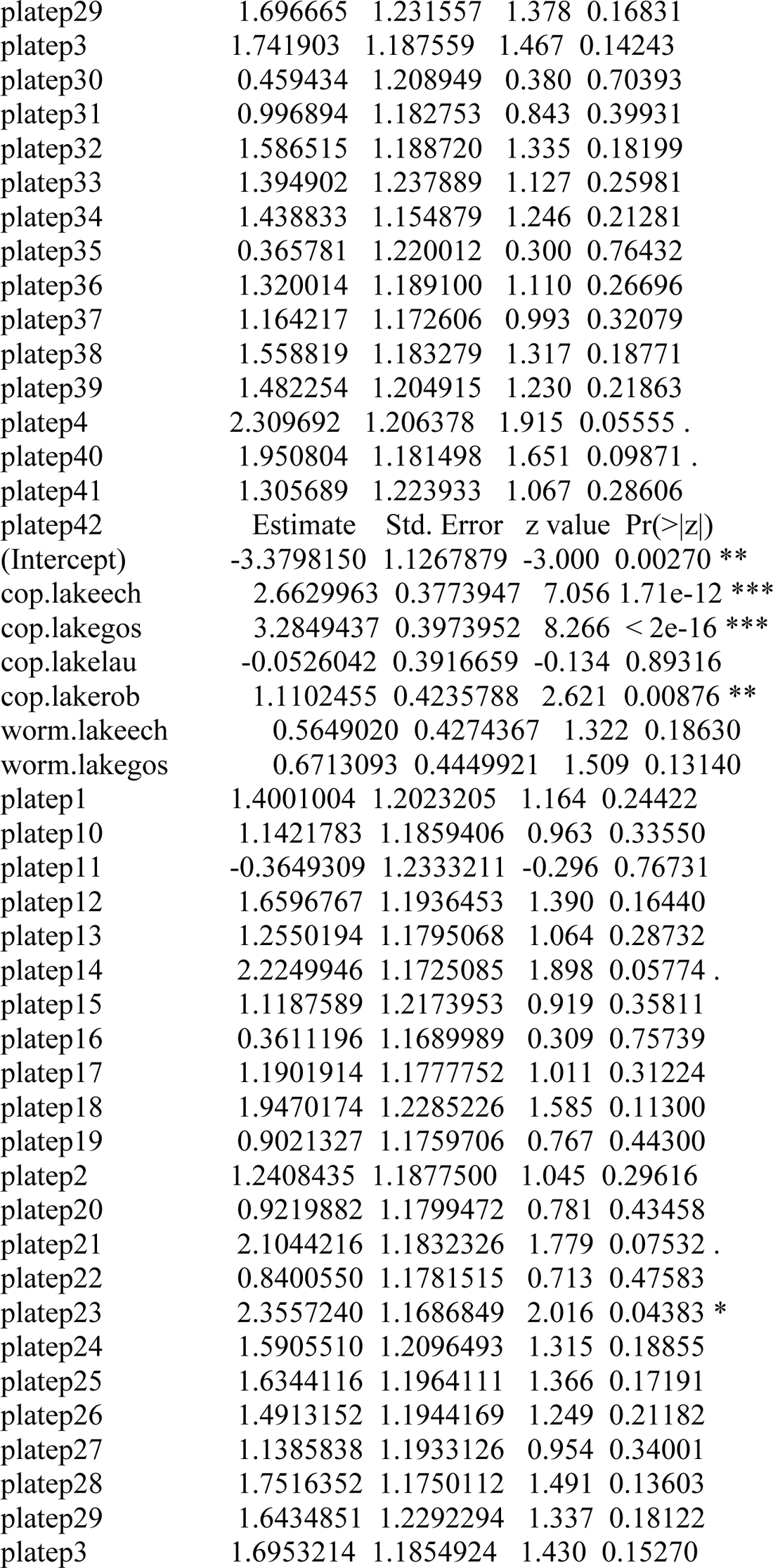
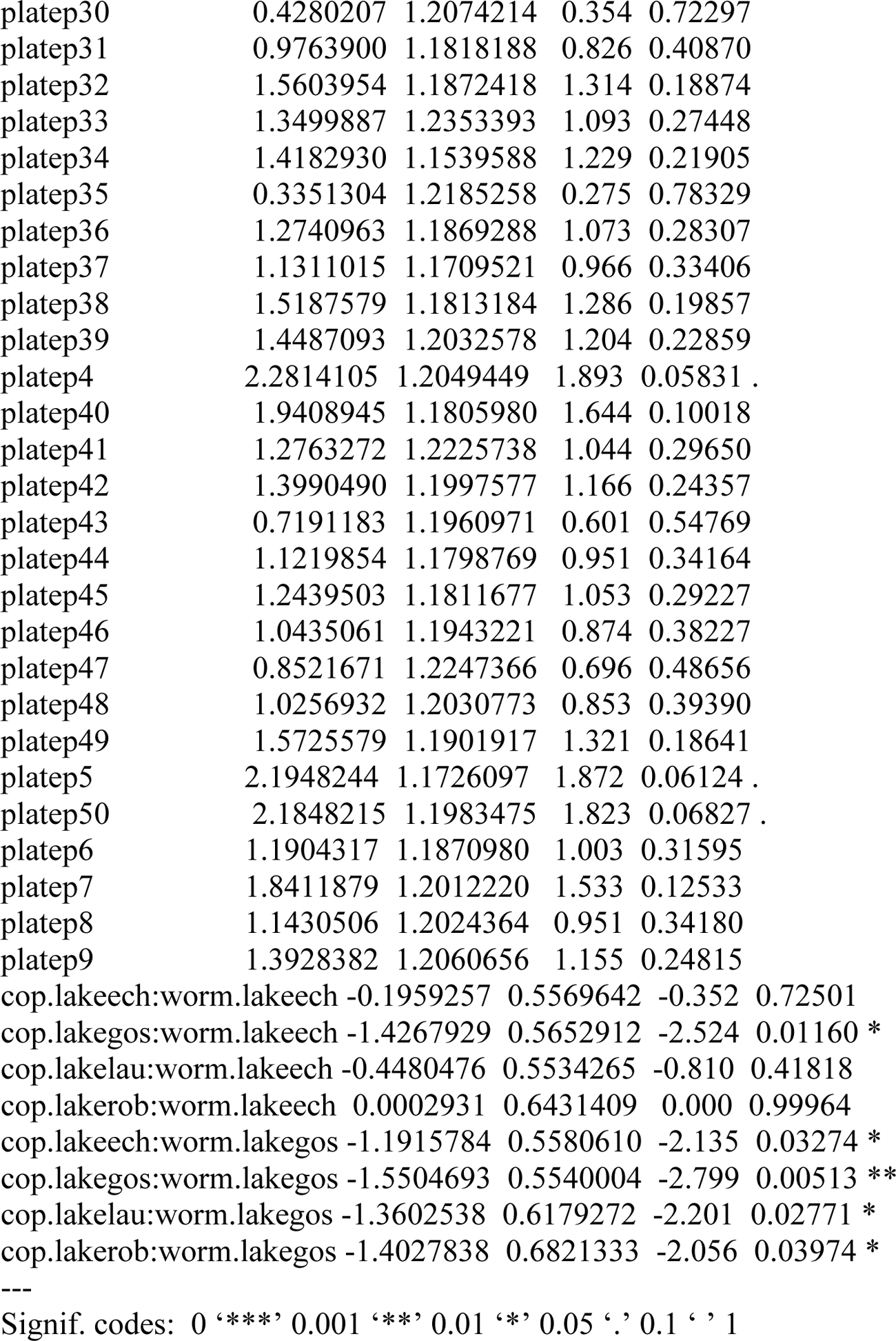

(Dispersion parameter for binomial family taken to be 1)

Null deviance: 2167.1 on 1621 degrees of freedom

Residual deviance: 1658.6 on 1557 degrees of freedom

AIC: 1788.6

Number of Fisher Scoring iterations: 5

> anova(model7, test = “LRT”)

Analysis of Deviance Table

Model: binomial, link: logit

Response: infected.yes.no

Terms added sequentially (first to last)

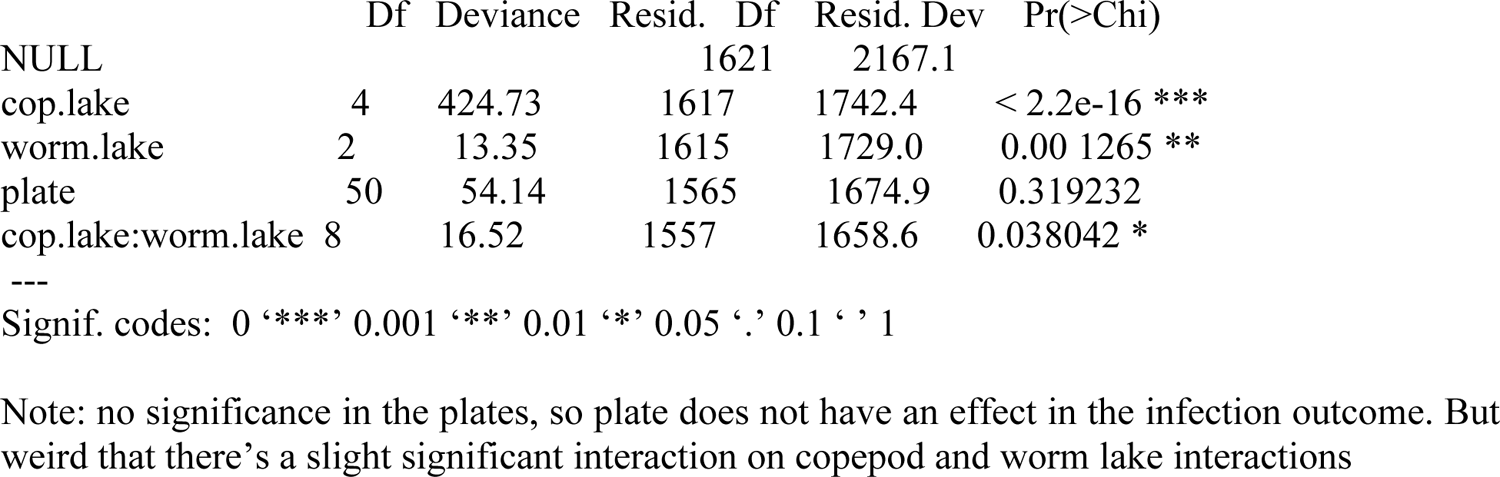

**5) testing for local adaptation**

> model8 <-glm(infected.yes.no ∼ cop.lake + worm.lake + native, + data=copepods, family=“binomial”)

> summary(model8)

Call:

glm(formula = infected.yes.no ∼ cop.lake + worm.lake + native, family = “binomial”, data = copepods)

Deviance Residuals:

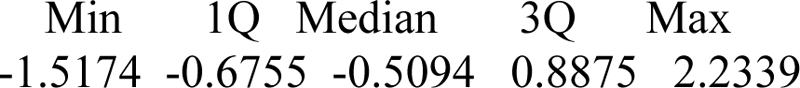

Coefficients:

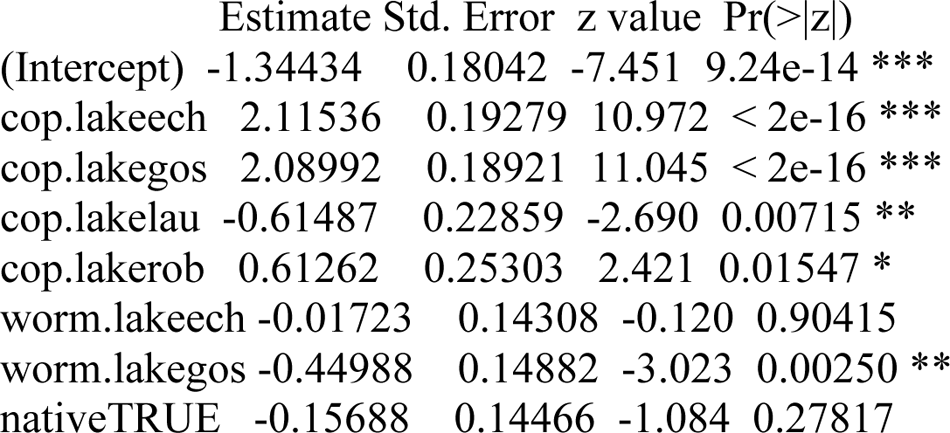
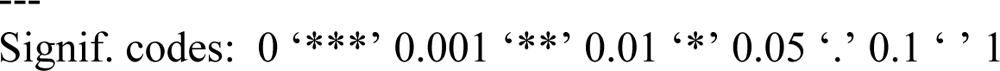

(Dispersion parameter for binomial family taken to be 1)

Null deviance: 2167.1 on 1621 degrees of freedom

Residual deviance: 1727.8 on 1614 degrees of freedom

AIC: 1743.8

Number of Fisher Scoring iterations: 4

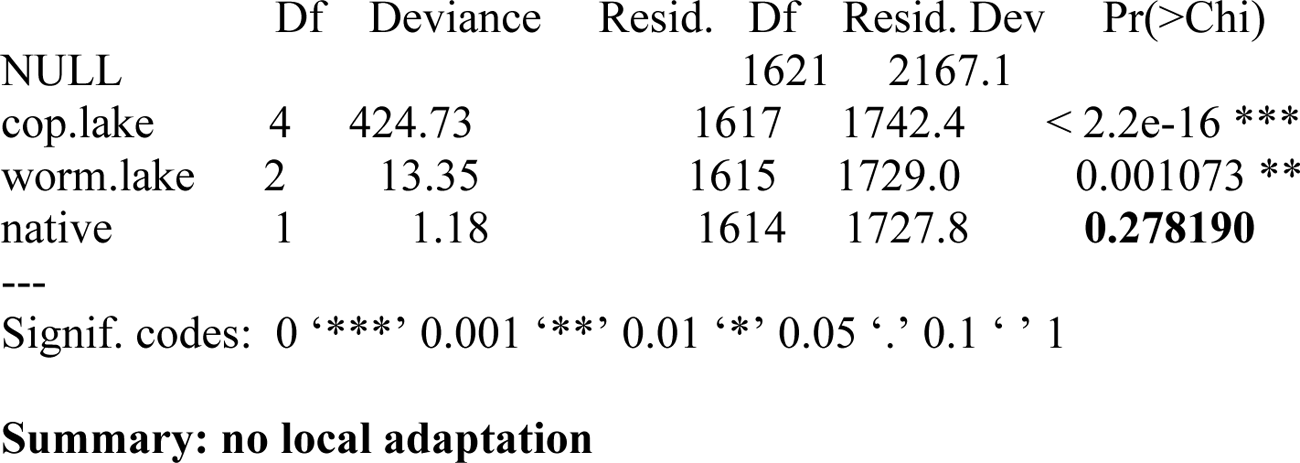

**6) testing for effect of worm family used: (summary answer after running the model below: tapeworm family does not matter)**

> model9 <-glm(infected.yes.no ∼ cop.lake + worm.lake + cop.lake*worm.lake + worm.fam,

+ data=copepods,family=“binomial”)

> summary(model9)

Call:

glm(formula = infected.yes.no ∼ cop.lake + worm.lake + cop.lake * worm.lake + worm.fam, family = “binomial”, data = copepods)

Deviance Residuals:

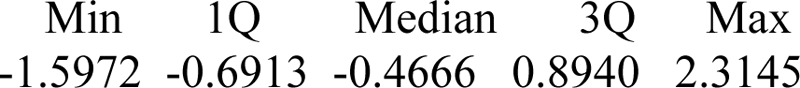

Coefficients: (2 not defined because of singularities)

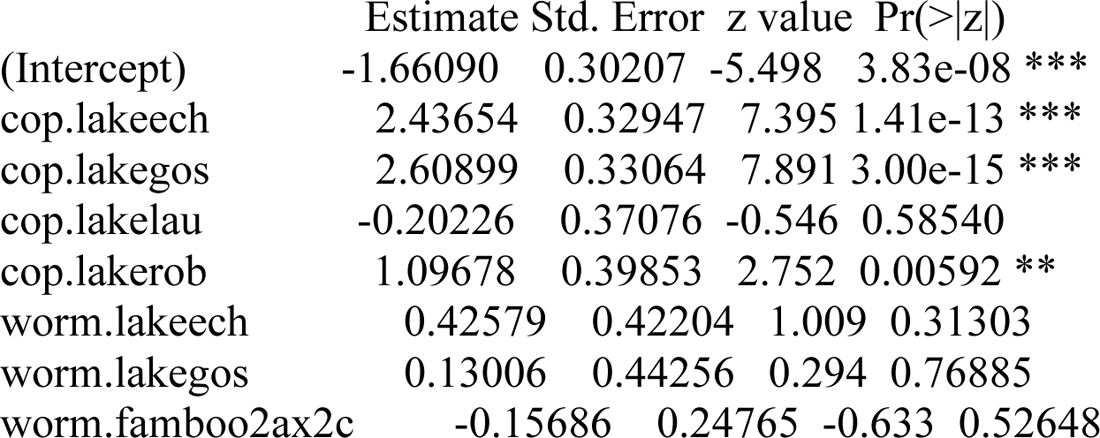
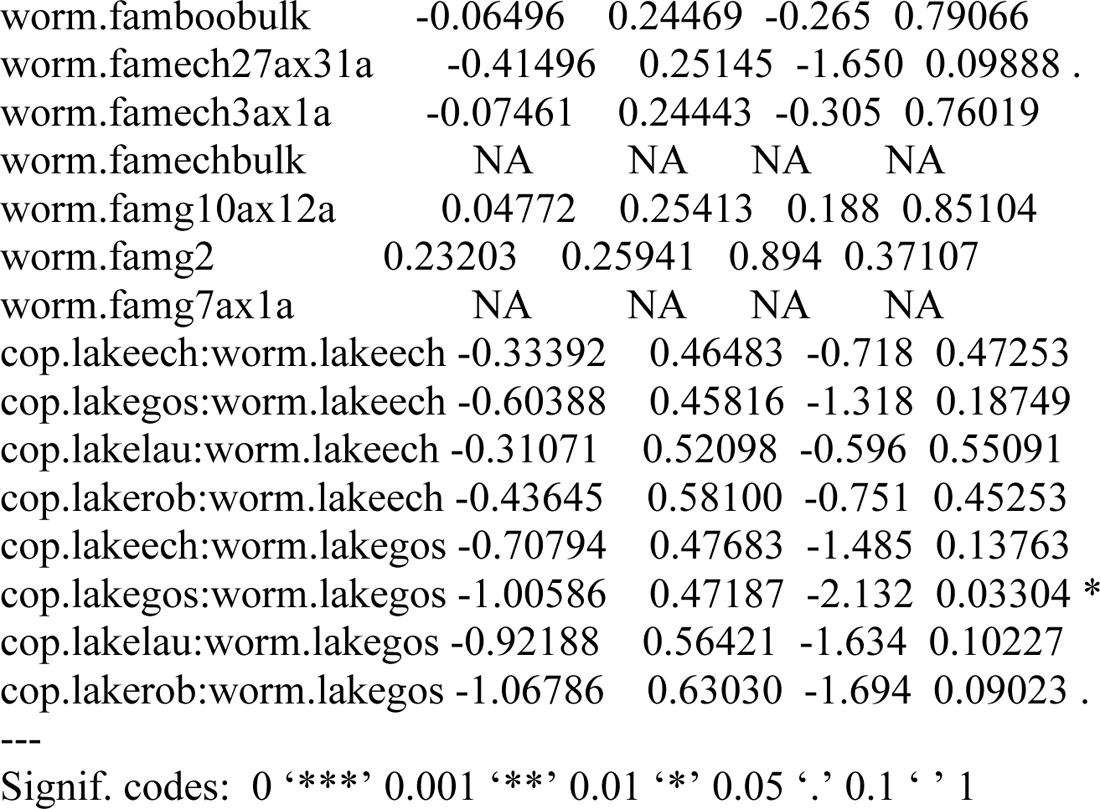

(Dispersion parameter for binomial family taken to be 1)

Null deviance: 2167.1 on 1621 degrees of freedom

Residual deviance: 1727.2 on 1599 degrees of freedom

AIC: 1773.2

Number of Fisher Scoring iterations: 13

> anova(model9, test = “LRT”)

Analysis of Deviance Table

Model: binomial, link: logit

Response: infected.yes.no

Terms added sequentially (first to last)

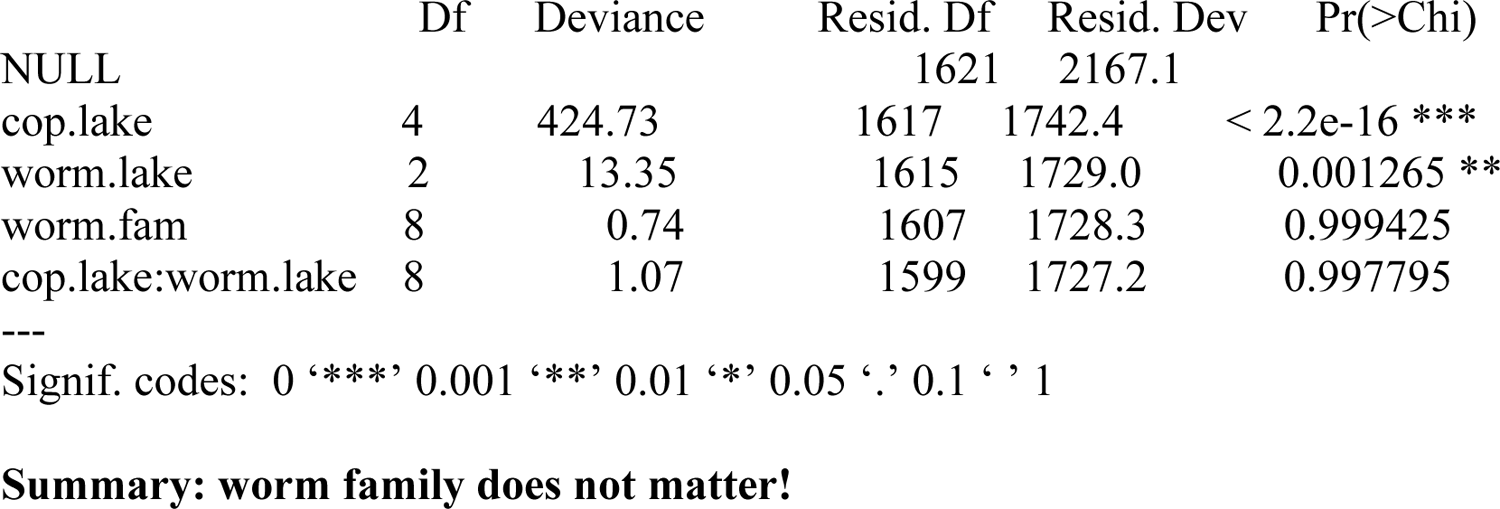

### Comparing all models on testing the prevalence of infection

> AIC(model1, model2, model3,model4, model5, model6, model7, model8, model9)

Note: according to lab-mate Christopher Peterson, it is fine to do AIC comparisons between GLM and GLMM models (need to ask him for the reference).

Below are the models sorted from best to worst: (the first number after each model name is the degrees of freedom followed by the Akaike Information Criterion (AIC) value:

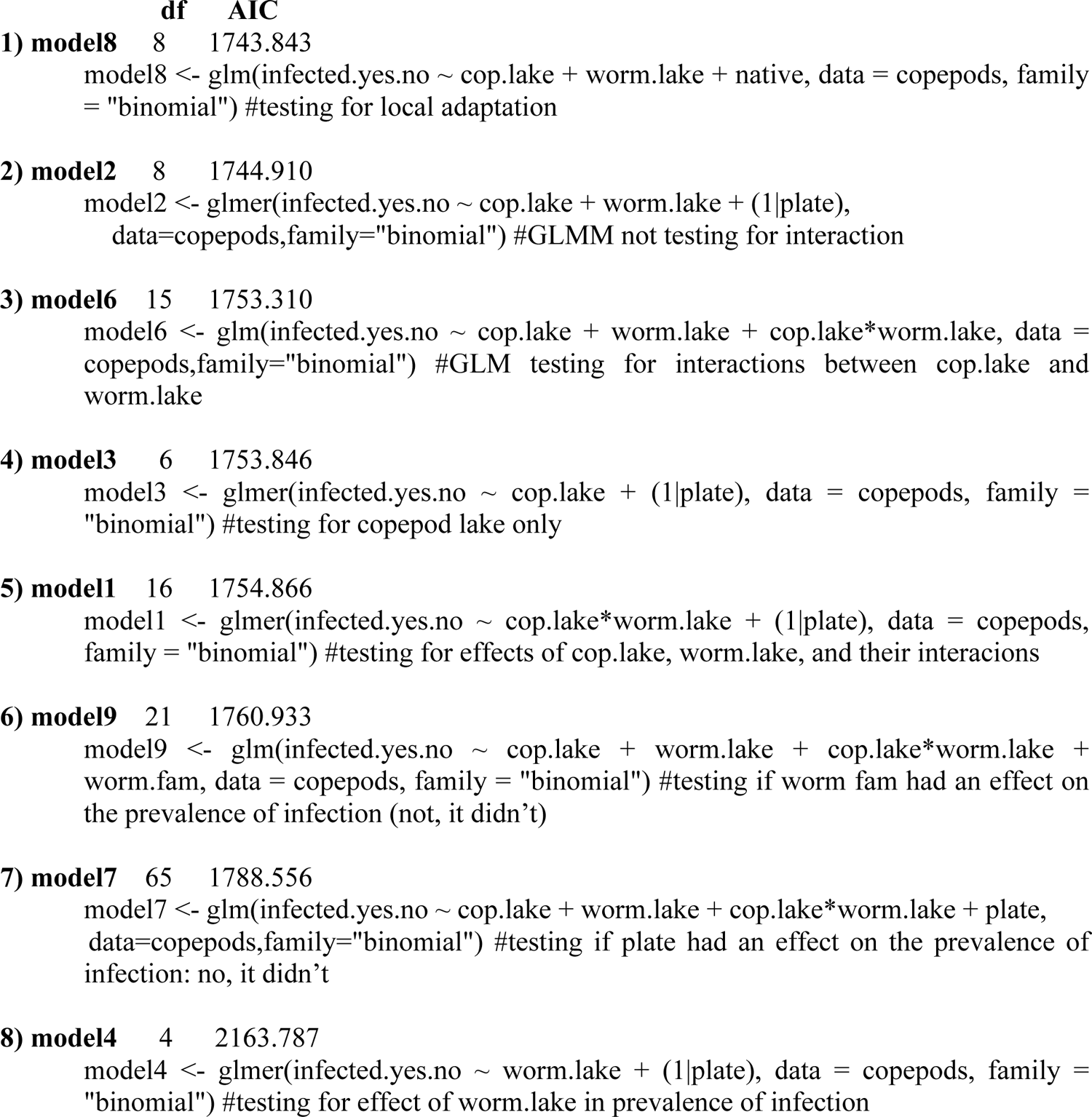
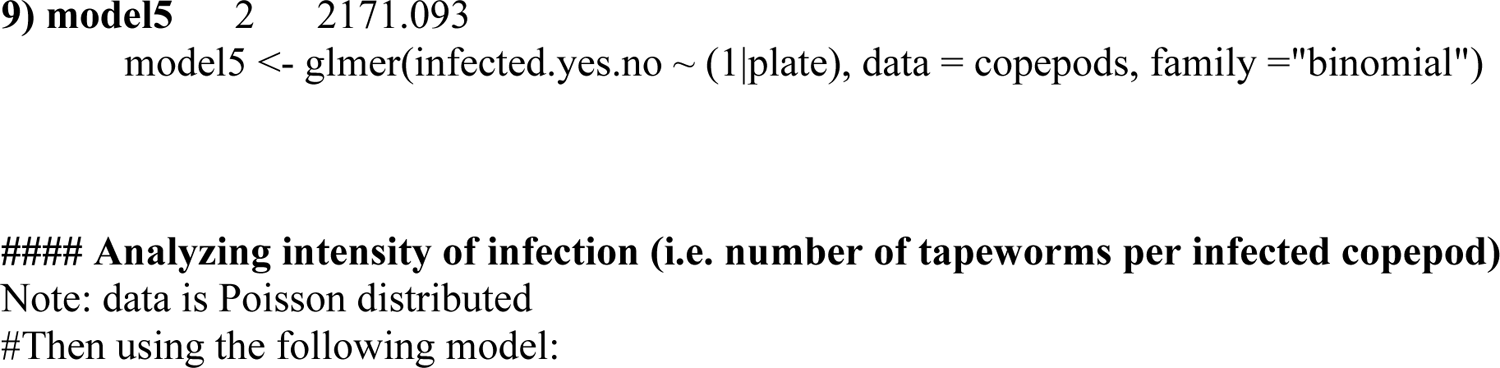

**7) Analyzing results on intensity including those not infected (i.e. number of worms >= 0)**

> model10 = glm (numb.worm ∼ cop.lake + worm.lake + cop.lake*worm.lake,

+ data=copepods,family=“poisson”)

> summary(model10)

Call:

glm(formula = numb.worm ∼ cop.lake + worm.lake + cop.lake * worm.lake, family = “poisson”, data = copepods)

Deviance Residuals:

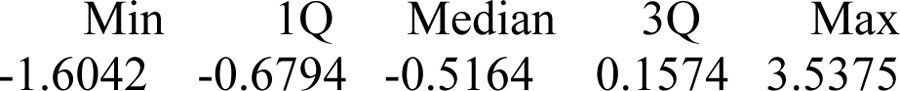

Coefficients:

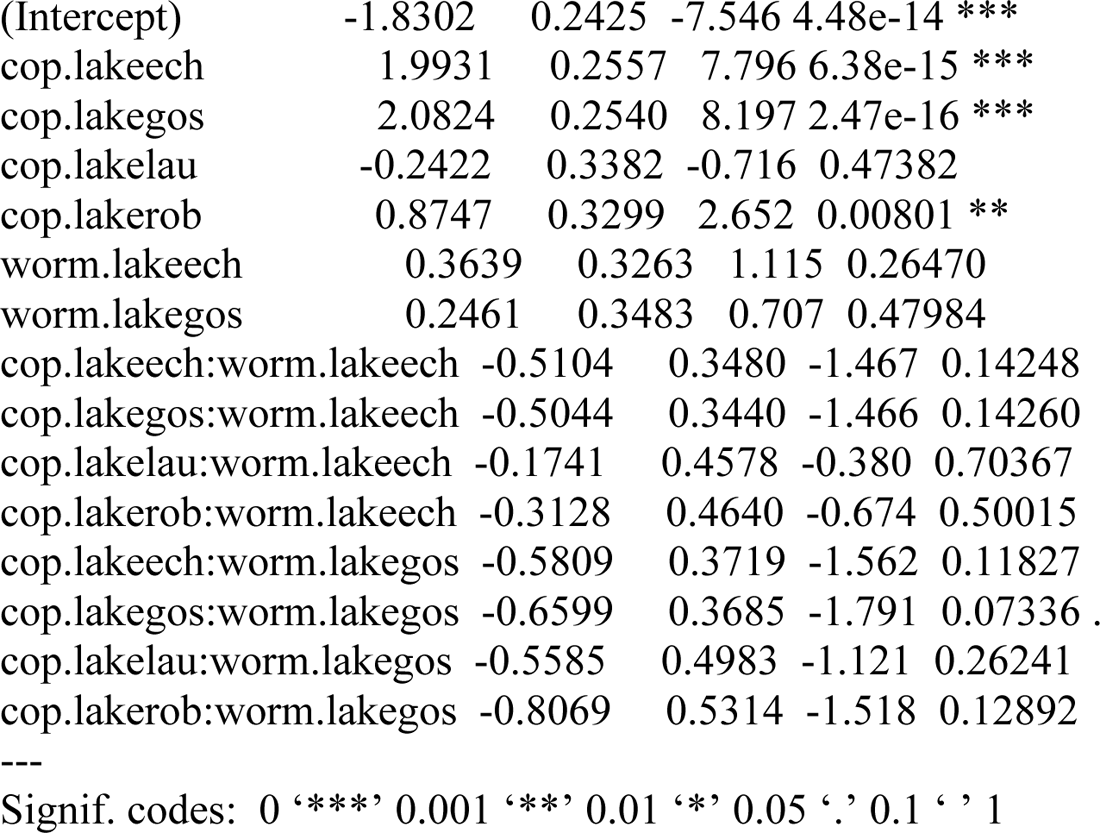

(Dispersion parameter for poisson family taken to be 1)

Null deviance: 2107.7 on 1621 degrees of freedom

Residual deviance: 1498.2 on 1607 degrees of freedom

AIC: 2959.8

Number of Fisher Scoring iterations: 6

> anova(model10, test = “LRT”)

Analysis of Deviance Table

Model: poisson, link: log

Response: numb.worm

Terms added sequentially (first to last)

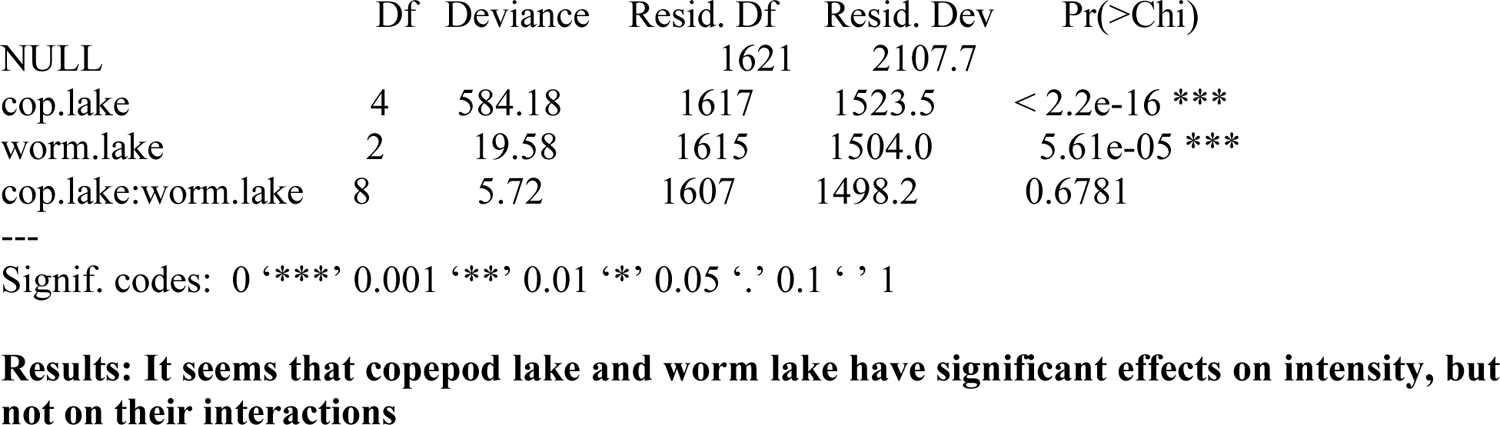

**8) What if I include a fixed variable in there, let’s say tapeworm family, and using “glmer” for GLMM**

> model11 <-glmer(numb.worm ∼ cop.lake*worm.lake + (1|worm.fam),

+ data=copepods,family=“poisson”)

> summary(model11)

Generalized linear mixed model fit by maximum likelihood (Laplace Approximation) [glmerMod] Family: poisson (log)

Formula: numb.worm ∼ cop.lake * worm.lake + (1 | worm.fam)

Data: copepods

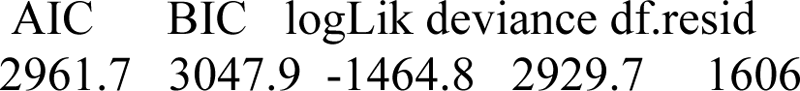

Scaled residuals:

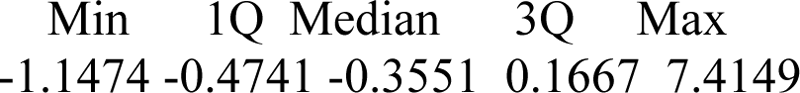

Random effects:

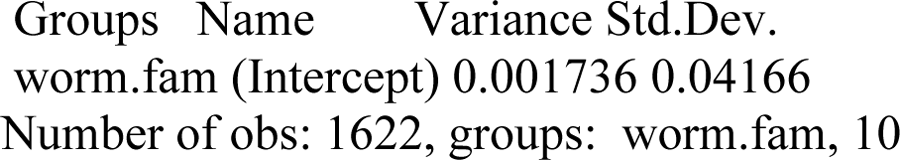

Fixed effects:

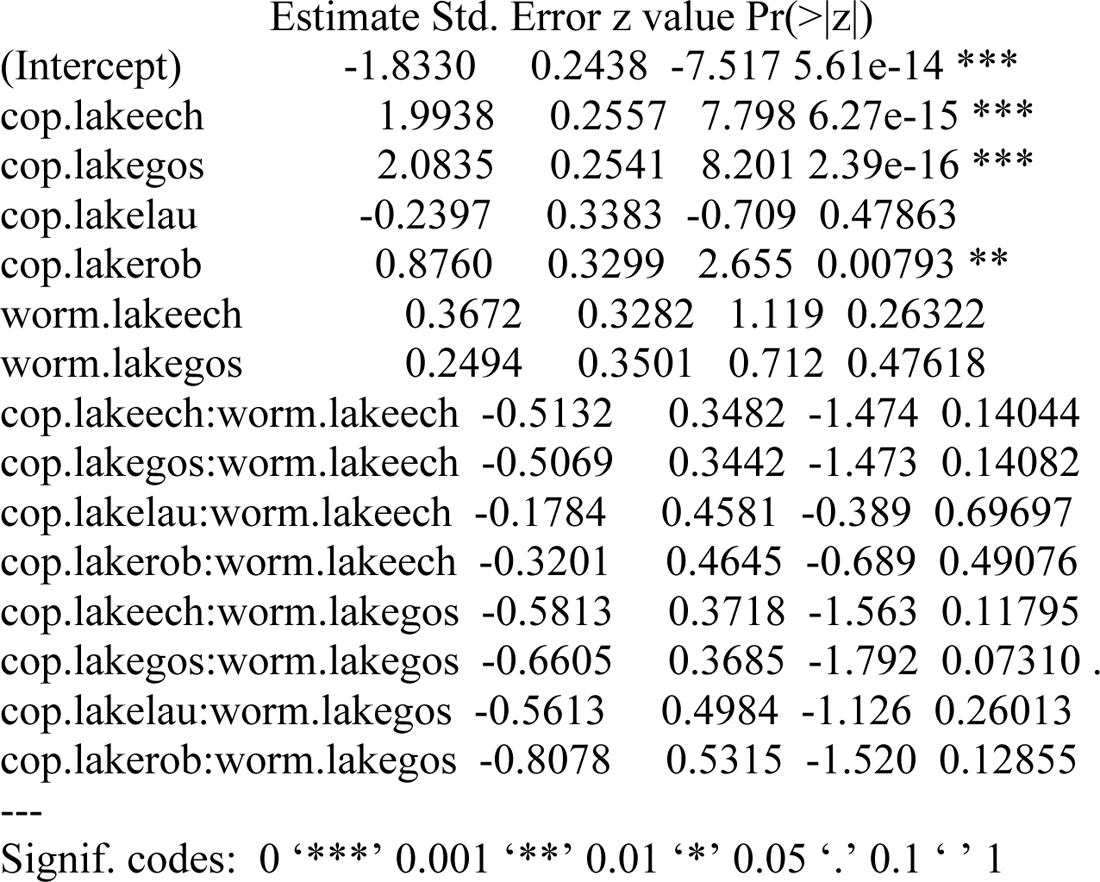

> AIC(model10,model11)

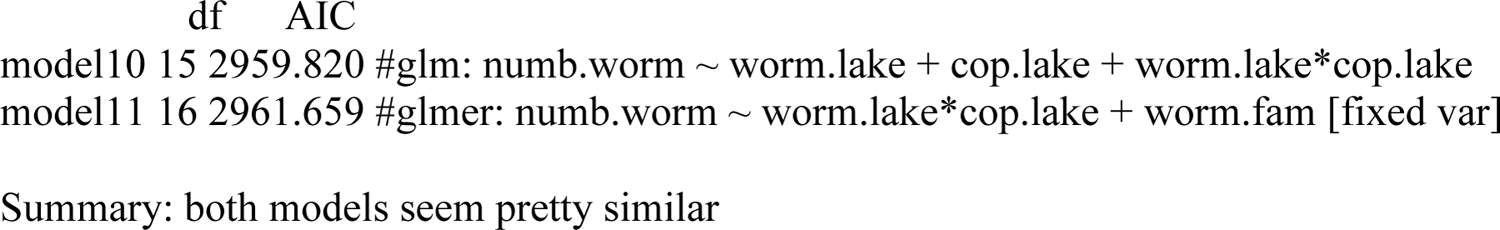

**9) what if I select only those copepods that got infected for the intensity analysis (as it should be)?**

preva = filter (copepods, numb.worm > 0) #using “filter” in “dplyer” R package to extract infected cops from dataset.

hist(preva$numb.worm, ylab = “# copepods”)

#Data is still Poisson distributed.

model12 = glm (numb.worm ∼ cop.lake + worm.lake + cop.lake*worm.lake, data = preva,family=“poisson”)

Deviance Residuals:

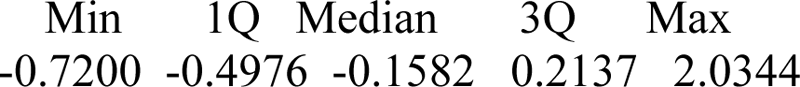

Coefficients:

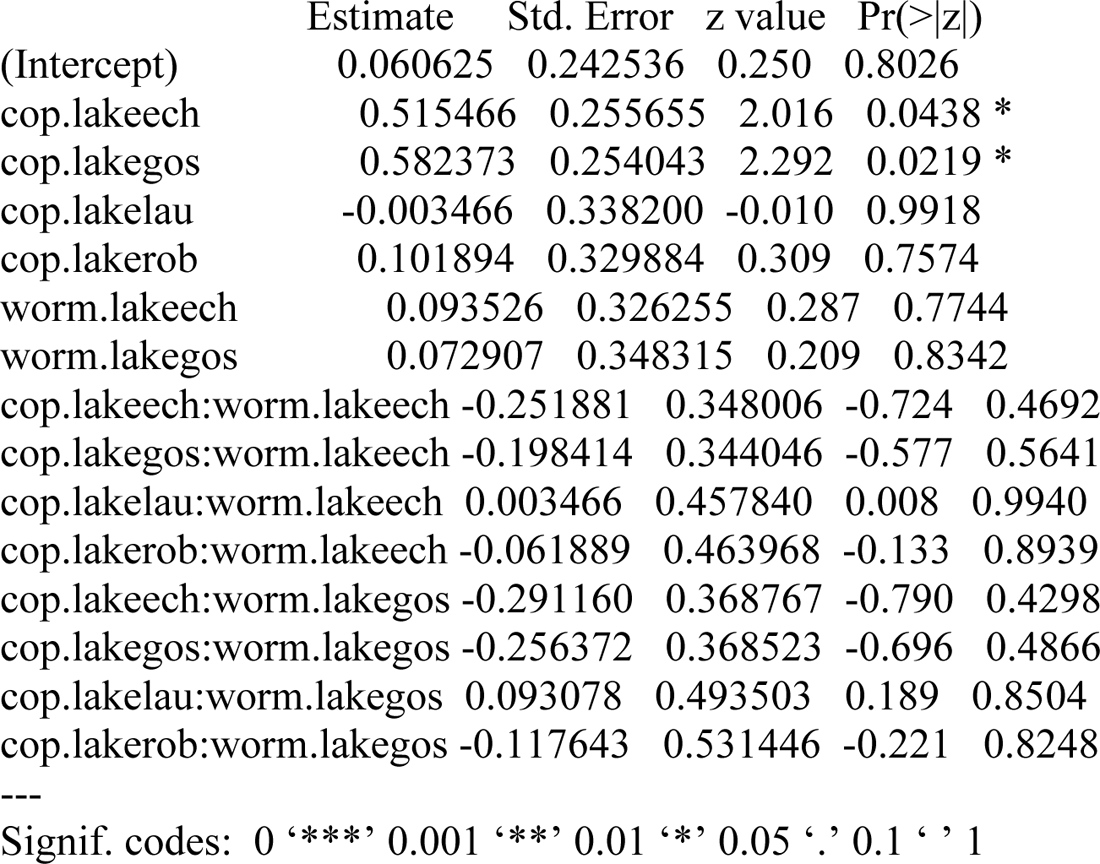

(Dispersion parameter for poisson family taken to be 1)

Null deviance: 238.68 on 621 degrees of freedom

Residual deviance: 211.90 on 607 degrees of freedom

AIC: 1673.5

Number of Fisher Scoring iterations: 4

> anova(model12, test = “LRT”)

Analysis of Deviance Table

Model: Poisson, link: log

Response: numb.worm

Terms added sequentially (first to last)

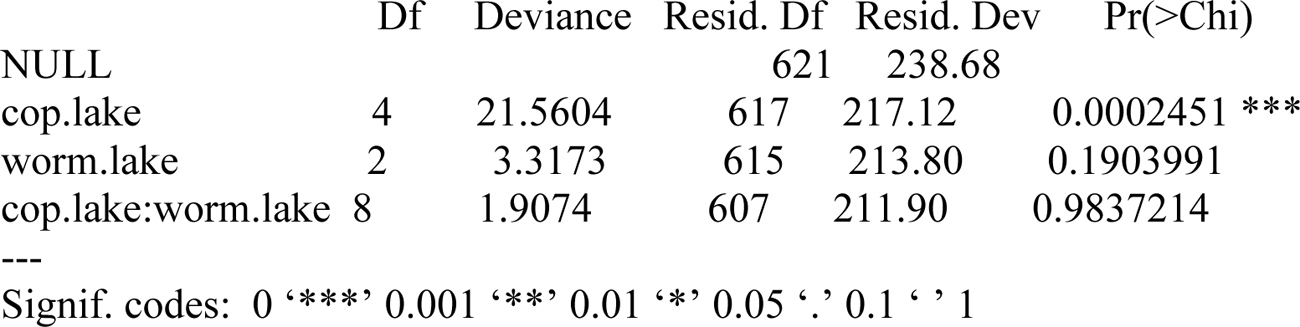
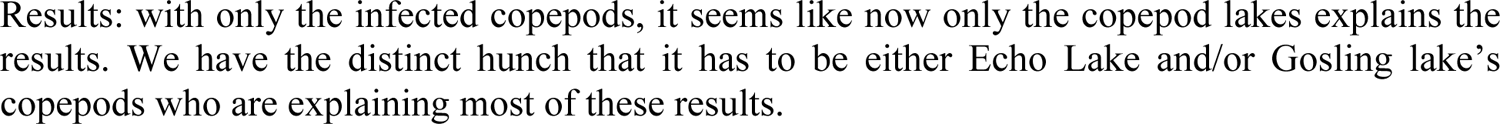

**10) What if we include a GLMM model using worm fam as the fix variable (only for the infected copepods; this is for intensity):**

Generalized linear mixed model fit by maximum likelihood (Laplace Approximation) model13 <-glmer(numb.worm ∼ cop.lake*worm.lake + (1|worm.fam), data = preva,family=“poisson”)

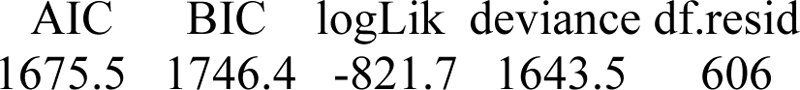

Scaled residuals:

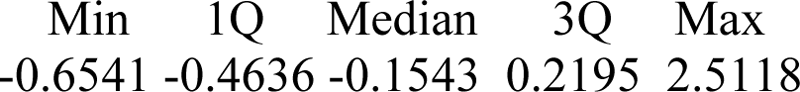

Random effects:

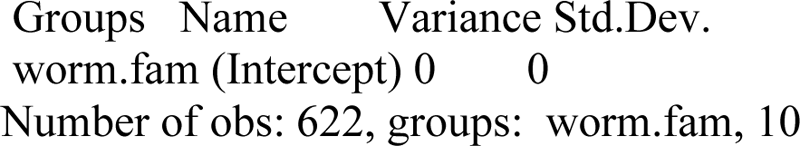

Fixed effects:

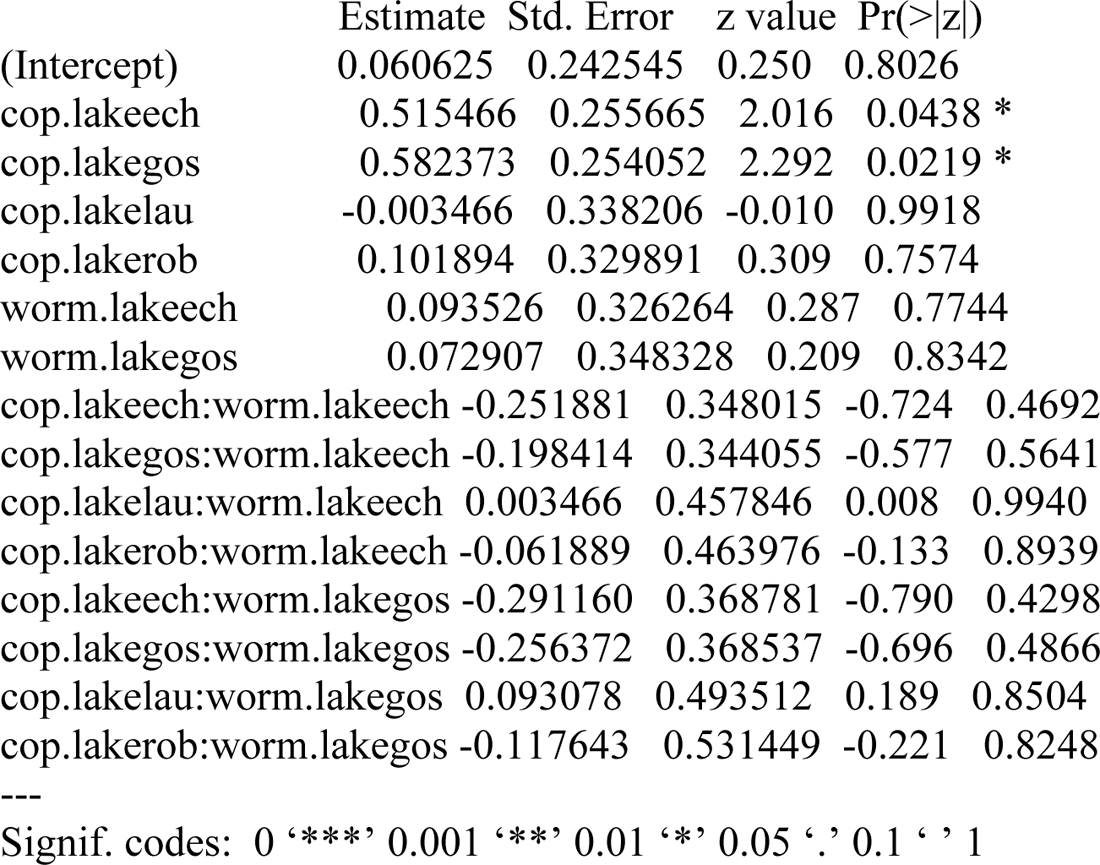

> AIC(model12,model13)

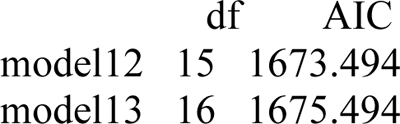
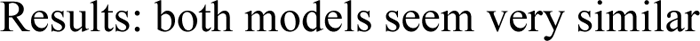

**11) Local adaption on intensity of infection levels (using only infected copepods for analyses):**

> summary(model16)

Call:

glm(formula = numb.worm ∼ cop.lake + worm.lake + native, family = “poisson”, data = preva)

Deviance Residuals:

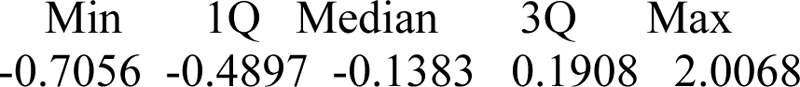

Coefficients:

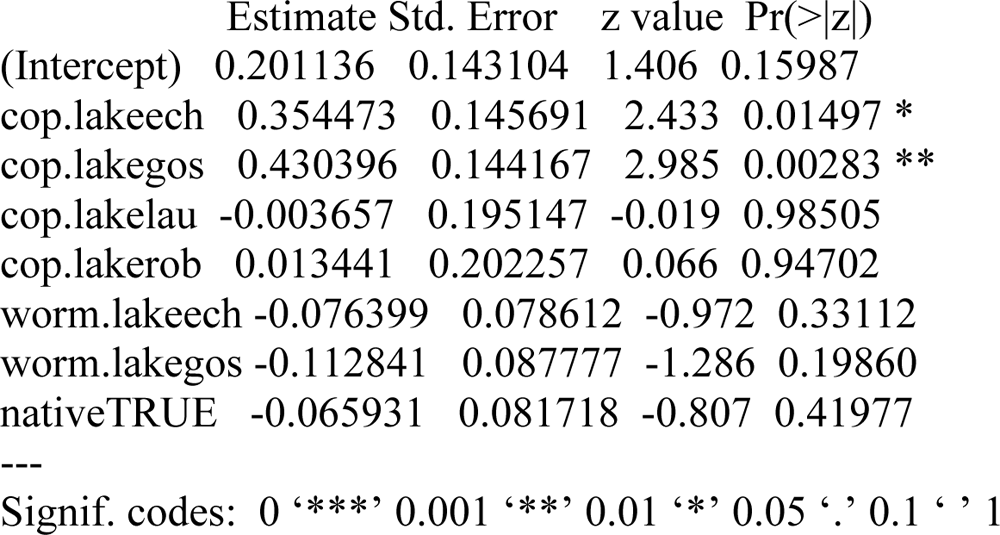

(Dispersion parameter for poisson family taken to be 1)

Null deviance: 238.68 on 621 degrees of freedom

Residual deviance: 213.15 on 614 degrees of freedom

AIC: 1660.7

> anova(model16, test = “LRT”)

Analysis of Deviance Table

Model: Poisson, link: log

Response: numb.worm

Terms added sequentially (first to last)

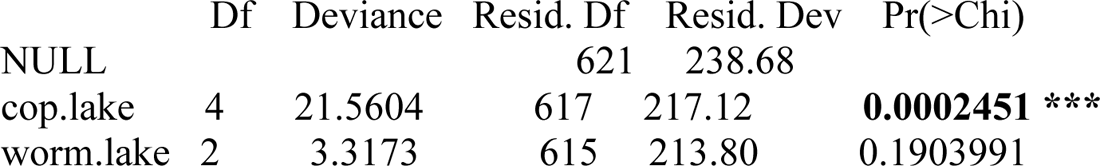
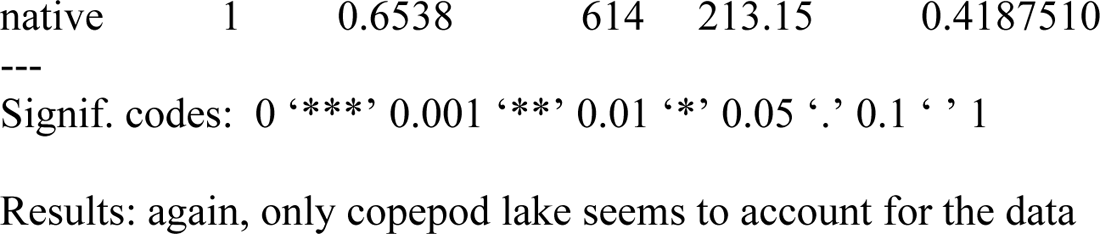

### The best models for intensity of infection (using only the data from infected copepods)

Below are the models sorted from best to worst: (the first number after each model name is the degrees of freedom followed by the Akaike Information Criterion (AIC) value:

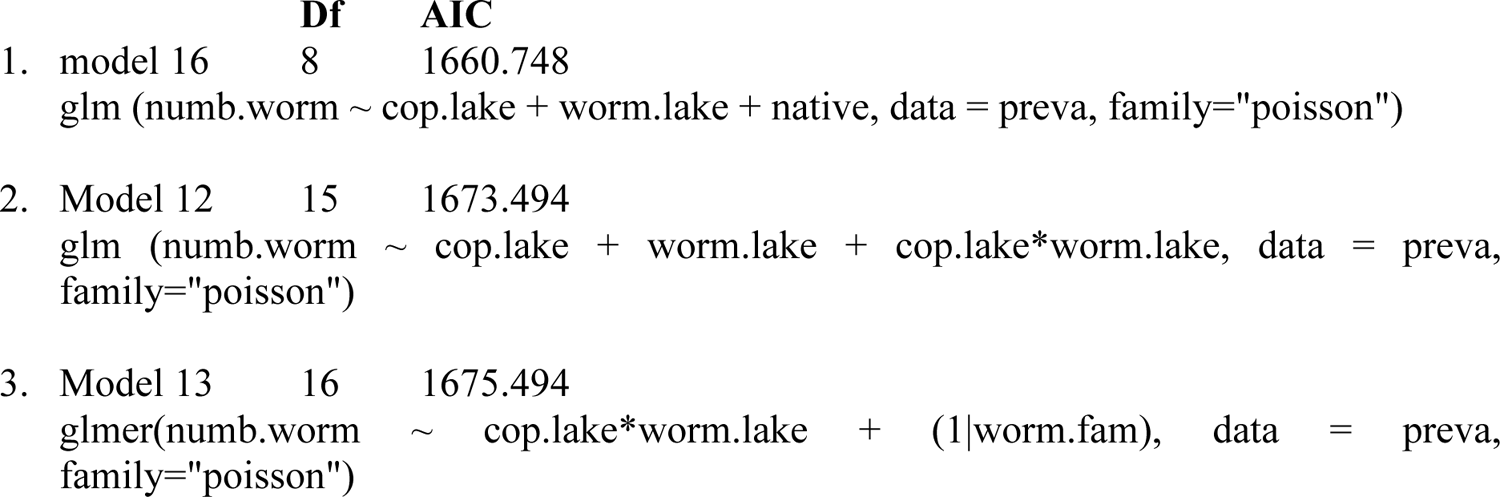

## References

1. Barber, I. 2013. Sticklebacks as model hosts in ecological and evolutionary parasitology. Trends in Parasitology, 29: 556–566.

2. Barber, I. and J. P. Scharsack. 2009. The three-spined stickleback-Schistocephalus solidus system: an experimental model for investigating host-parasite interactions in fish. Parasitology, 137: 411–424.

3. Benesh, D. P. 2010. What are the evolutionary constraints on larval growth in a trophically transmitted parasite? Oecologia, 162: 599–608.

4. Bürkner P. C. 2018. Advanced Bayesian Multilevel Modeling with the R Package brms. The R Journal. 10(1), 395–411. doi.org/10.32614/RJ-2018-017

5. Caldera, E.J. and Bolnick, D. I. 2008. Effects of colonization history and landscape structure on genetic variation within and among threespine stickleback (Gasterosteus aculeatus) populations in a single watershed. Evolutionary Ecology Research, 2008: 575–598.

6. Dubinina M. N. Tapeworms (Cestoda, Ligulidae) of the fauna of the USSR (Translated from Russian). New Delhi: Amerind Publishing Co. Pvt. Ltd., 1980.

7. Gandon, S., and Nuismer, S.L. 2009. Interactions between Genetic Drift, Gene Flow, and Selection Mosaics Drive Parasite Local Adaptation. The American Naturalis, 173: 212–224.

8. Greischar, M.A. and Koskella, B., 2007. A synthesis of experimental work on parasite local adaptation. Ecology letters, 10:418–434.

9. Hafer, N., 2018. Differences between populations in host manipulation by the tapeworm Schistocephalus solidus–is there local adaptation? Parasitology, 145(6), pp.762–769.

10. Haney, J.F., et al. “An-Image-based Key to the Zooplankton of North America” version 5.0 released 2013. University of New Hampshire Center for Freshwater Biology <cfb.unh.edu> 1 Oct 2019.

11. Hoberg, E. P., Henny, C. J., Hedstrom, O. R. and Grove, R. A. 1997. Intestinal helminths of river otters (Lutra canadensis) from the Pacific Northwest. Journal of Parasitology, 83: 105–110.

12. Hoeksema, J.D. and Forde, S.E., 2008. A meta-analysis of factors affecting local adaptation between interacting species. The American Naturalist, 171(3), pp.275–290.

13. Kalbe, M., Eizaguirre, C., Scharsack, J.P. and Jakobsen, P.J., 2016. Reciprocal cross infection of sticklebacks with the diphyllobothriidean cestode Schistocephalus solidus reveals consistent population differences in parasite growth and host resistance. Parasites & vectors, 9(1), pp.1–12.

14. Lenormand T. 2002. Gene flow and the limit to natural selection. Trends in Ecology and Evolution, 17: 183–189

15. Lively, C.M., M.F. Dybdahl, J. Jokela, E. Osnas, L.F. Delph. 2004. Host sex and local adaptation by parasites in a snail-trematode interaction. American Naturalist, 164:S6–S18.

16. Mazé-Guilmo, E., Blanchet, S., McCoy, K.D. and Loot, G., 2016. Host dispersal as the driver of parasite genetic structure: a paradigm lost? Ecology Letters, 19(3), pp.336–347.

17. Morgan, A.D., Gandon, S., and Buckling, A. 2005. The effect of migration on local adaptation in a coevolving host–parasite system. Nature, 437:253–256.

18. Noble, E. R., Noble, G. A., Schad, G. A., and MacInnes, A. J. 1989. Parasitology: The Biology of Animal Parasites, 6th ed. Lea & Febiger, Philadelphia

19. Poulin, R., 2007. Evolutionary ecology of parasites. Princeton University Press, New Jersey.

20. R Core Team. 2018. R: A language and environment for statistical computing. R Foundation for Statistical Computing, Vienna, Austria. https://www.R-project.org/.

21. Schmid-Hempel, P., 2011. Evolutionary parasitology. Oxford University Press, Oxford, UK.

22. Smyth, J. D. 1990. In vitro Cultivation of Parasitic Helminths. CRC Press, Boca Raton, FL.

23. Weber, J. N., M. Kalbe, K. C. Shim, N. I. Erin, N. C. Steinel, L. Ma, and D. I. Bolnick. 2017a. Resist globally, infect locally: a transcontinental test of adaptation by stickleback and their tapeworm parasite. The American Naturalist, 189: 43–57.

24. Weber, J. N., N. C. Steinel, K. C. Shim, and D. I. Bolnick. 2017b. Recent evolution of extreme cestode growth suppression by a vertebrate host. Proceedings of the National Academy of Sciences, 114: 6575–6581.

25. Wedekind, C. 1997. The infectivity, growth, and virulence of the cestode Schistocephalus solidus in its first intermediate host, the copepod Macrocyclops albidus. Parasitology, 115: 317–324.

26. Yao, Y., A. Vehtari, D. Simpson, and A. Gelman. 2018. Using Stacking to Average Bayesian Predictive Distributions (with Discussion). Bayesian Analysis. 13 (3): 917–1007

